# von Willebrand Factor D and EGF Domains is an evolutionarily conserved and required feature of blastemas capable of multi-tissue appendage regeneration

**DOI:** 10.1101/842948

**Authors:** N.D. Leigh, S. Sessa, A.C. Dragalzew, D. Payzin-Dogru, J.F. Sousa, A.N. Aggouras, K. Johnson, G.S. Dunlap, B.J. Haas, M. Levin, I. Schneider, J.L. Whited

## Abstract

Regenerative ability varies tremendously across species. A common feature of regeneration of appendages such as limbs, fins, antlers, and tails is the formation of a blastema--a transient structure that houses a pool of progenitor cells that regenerate the missing tissue. We have identified the expression of *von Willebrand Factor D and EGF Domains* (*vwde*) as a common feature of blastemas capable of regenerating limbs and fins in a variety of highly regenerative species. Further, *vwde* expression is tightly linked to the ability to regenerate appendages. Functional experiments demonstrate a requirement for *vwde* in regeneration and indicate that Vwde is a potent mitogen in the blastema. These data identify a key role for *vwde* in regenerating blastemas and underscore the power of an evolutionarily-informed approach for identifying conserved genetic components of regeneration.

## Introduction

The underlying reasons why some animals have the ability to regenerate complex structures, while others cannot, remains an important and open question. This knowledge gap has led to intense study of how regeneration-competent species are able to perform complex multi-tissue regeneration, with a particular focus on the ability to regenerate paired appendages, such as limbs and fins. However, this has long been a pursuit without an understanding of whether this ability was present when paired appendages first evolved or was acquired by certain phylogenetic lineages (e.g. urodele amphibians).

Recent work regarding the evolutionary origins of regenerative capacity has indicated that the ability to regenerate paired appendages is an inherited feature of the fin-to-limb transition [1–4]. Evidence found in the fossil record [3,4], functional studies across species [2], and comparisons of gene expression profiles of regenerating tissue [1,2] support the notion that paired appendage regeneration is a feature lost by certain lineages and was not a newly derived capacity in highly regenerative lineages. This indicates that the amniote lineage (which includes humans) has lost regenerative tendencies in appendages over evolutionary time. Therefore, the ability to stimulate regeneration in non-regenerative species, potentially in a therapeutic context, may require the re-initiation of a core, evolutionary conserved program.

All species that are able to regenerate appendages share a conserved trait: the ability to form a blastema. The blastema is the morphological structure that forms at the amputation plane and houses the progenitor cells responsible for regeneration. Recent efforts have focused on elucidating the molecular definition of the blastema, with many of these efforts aimed at the axolotl limb blastema due to the ease of tissue acquisition and the ability to perform experimentation in the lab [5–13]. These studies provide a wealth of information about transcriptomic changes over time, cell types, and blastema-enriched genes. More recently, sequencing efforts of non-model species have allowed for comparisons to the axolotl limb blastema and indicate a core molecular signature that is shared between the blastemas of distantly related species [1,2]. Due to these similarities and the common evolutionary origin of limb regeneration capacity, we can use an evolutionarily-informed approach to understand what constitutes a blastema and for identifying core features required for regeneration.

A recent approach to identify the unique gene expression in the axolotl limb blastema compared blastema gene expression to a variety of homeostatic and embryonic tissues and identified over 150 blastema-enriched genes [11]. These blastema-enriched genes may help to explain the unique functions of the blastema, but the question remains as to whether these genes represent a core program or are functionally required for regeneration. One of the most blastema-enriched genes in this dataset was *von Willebrand Factor D and EGF Domains* (*vwde*), which to date has not been functionally studied in any context.

We decided to apply an evolutionary framework to determine if *vwde* fit the description of an evolutionary-conserved, blastema-enriched gene and if such an approach may help to identify genes required for regeneration. We found that *vwde* expression is a common feature of both fin and limb blastemas and was highly enriched in regenerating appendages as compared to pre-amputation intact appendages. In addition, using the natural regeneration-competent and regeneration-refractory periods during *Xenopus laevis* development, we observed that *vwde* expression was tightly linked to the regeneration-component environment. This suggests that *vwde* may be a critical factor in the regenerative niche. Finally, we found that *vwde* is functionally required for axolotl limb regeneration, with transient knockdown of protein levels resulting in aberrant regeneration. These data suggest that an evolutionarily-informed approach can help to prioritize target genes and that genes that are blastema-enriched across different species may prove to be critical factors in the ability to regenerate appendages.

## Results

With the goal of identifying genes enriched to the regenerating blastema, a tissue-mapped axolotl transcriptome was recently published [11]. Of particular interest are blastema-enriched genes with high expression in the blastema and relative low expression in all other tissues sampled. We found that *vwde* was highly enriched to the axolotl limb blastema (Figure 1A). This analysis, however, was limited to one time point, the medium-bud blastema, and it did not provide spatial information about the expression of *vwde* across the regenerating limb. To understand the spatial and temporal regulation of *vwde*, we performed RNA *in situ* hybridizations over a time course of axolotl limb regeneration. We found *vwde* expression to be tightly tied to the presence of a blastema and expressed exclusively in the blastema and not the overlying wound epidermis (Figure 1B-E). Thus, *vwde* fits the description of an axolotl limb blastema-enriched gene and we were interested in pursuing whether *vwde* may be a core component of blastemas able to regenerate appendages.

**Figure 1:**
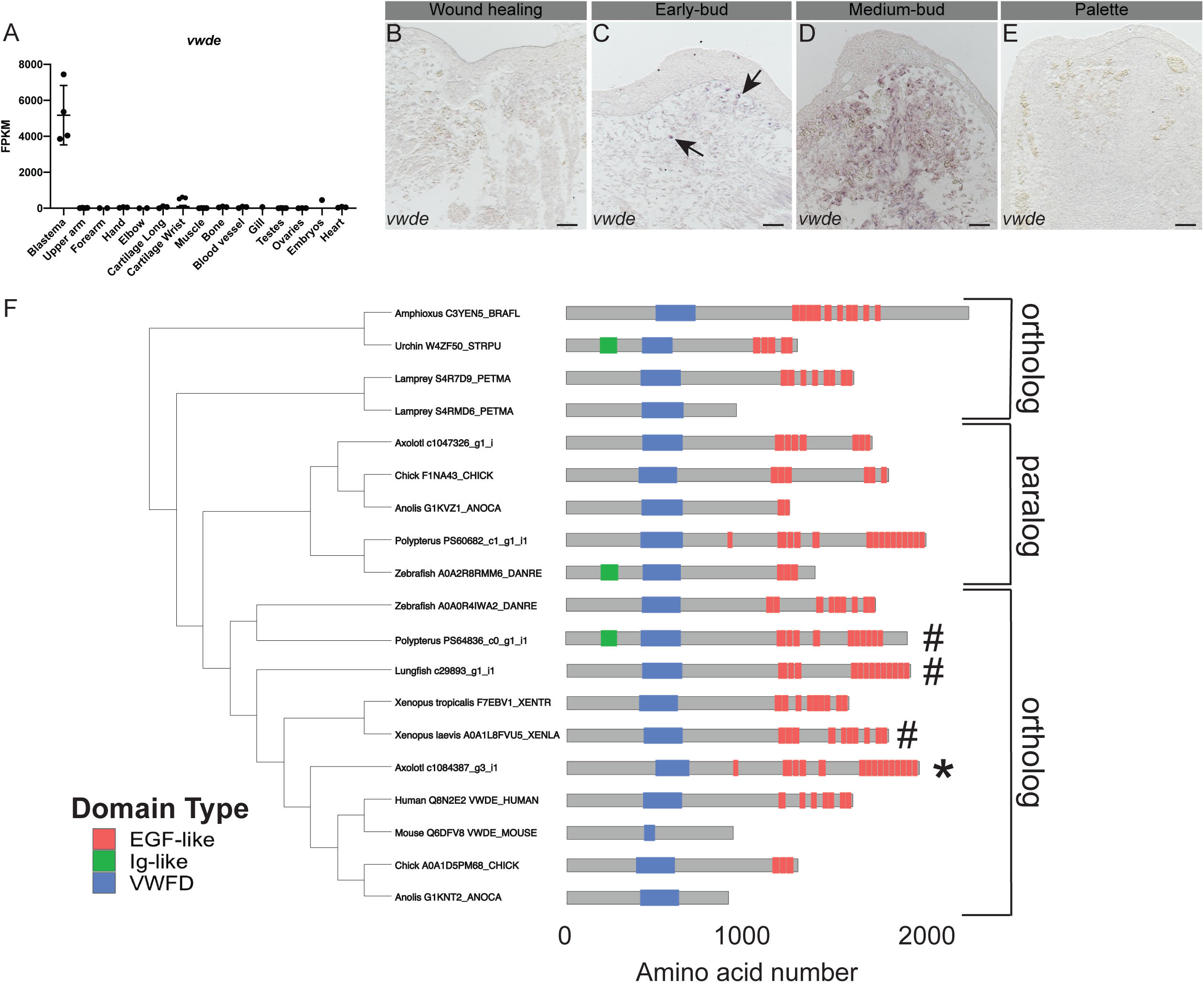
*von Willebrand Factor D and EGF-like Domains* (*vwde*) is a blastema-enriched gene that is found across deuterostomes. (A) *vwde* (contig c1084387_g3_i1) expression in FPKM across tissues sampled from Bryant et al. Cell Reports 2017. Proximal and distal blastema samples are combined. (B-E) RNA *in situ* hybridization for *vwde* at (B) wound healing, (C) early-bud blastema, (D) medium-bud blastema, and (E) palette stage regenerating limbs. Black arrows indicate *vwde* expression, scale bar is 100 µm. (F) OrthoFinder 2.0 phylogeny with corresponding protein domain structure for putative Vwde orthologs. Protein domain pictures were generated with drawProteins [38]. Species and Uniprot ID, transcriptome contig number, or Ensembl ID are included. *Polypterus vwde* contained multiple splice isoforms and the closest match to axolotl Vwde is shown here. Axolotl Vwde is denoted with (*) and other species Vwde that are described in this manuscript are marked with (#). Orthologs to the Vwde studied in this work are indicated with brackets, paralog is also denoted with brackets.

We next sought to determine if *vwde* was present in a selection of deuterostomes, including species with various regenerative abilities. Using a comparative genomics approach [14], we found *vwde* to have orthologs across deuterostomes (though no ortholog was detected in *Ciona intestinalis*), as well as a non-blastema-enriched paralogous gene in axolotl and other species (Figure 1F, Supplementary Figure 1). We compared axolotl VWDE to proteins from other species, and we found putative orthologs harboring predicted von Willebrand Factor D domains and EGF-like domains. The number of EGF domains may be more variable across species. However, since this gene has not yet been studied in-depth in any species, additional experimental work may be required to fully characterize the expressed transcripts and proteins for individual species. Using these identified orthologs, we moved forward to ask whether *vwde* was a blastema-enriched gene during paired fin regeneration.

We explored the possibility that *vwde* could be a common feature of blastemas responsible for regenerating paired fins, which share a deep homology with limbs [15], and likely share an inherited gene regulatory program for regeneration [1,2]. We chose two highly regenerative, but distantly related, fish species to determine if *vwde* expression was a conserved feature of blastemas capable of regenerating paired appendages. These include a species in the sister group to tetrapods, the Lungfish (*Lepidosiren paradoxa*), which is a lobe-finned fish, and *Polypterus senegalus*, a ray-finned fish that is capable of regenerating after amputation through skeletal elements that develop by endochondral ossification. We first inspected publicly available transcriptome datasets of lungfish and *Polypterus* regenerating fins for the *vwde* orthologs we previously identified (Figure 1F). The lungfish LG29893_g1_i1 contig was upregulated in blastemas 21 days post-amputation (dpa) relative to uninjured fins [1], and the *Polypterus* PS64836c0_g1_i1 contig was upregulated in 9 dpa blastemas relative to uninjured fins [2]. Assessment of expression levels via qPCR at various regeneration stages showed an upregulation of *vwde* coinciding with blastema formation during lungfish fin regeneration (Supplemental Figure 2A).

A similar pattern was seen for *Polypterus* fin regeneration, with expression reaching highest levels at 5 dpa (Supplemental Figure 2B). Next, we assessed the spatial pattern of *vwde* in histological sections of regenerating fins. Lungfish 21 dpa blastemas show distal mesenchymal expression of *vwde* (Figure 2A). In 5 dpa *Polypterus* blastemas, expression is observed distal to the amputation plane in mesenchymal cells but also in the epithelium, suggesting that *Polpyterus* may use *vwde* in both compartments (Figure 2C). *In situ* hybridization with control sense probes did not yield specific signal (Supplemental Figure 2C-D). Histologically, these samples are similar to the medium-bud blastema time point in which we identified *vwde* in the axolotl limb (Figure 2B, 2D). Together, these data indicate that *vwde* is expressed in regenerating fins and limbs and that *vwde* expression is a conserved feature of blastemas.

**Figure 2:**
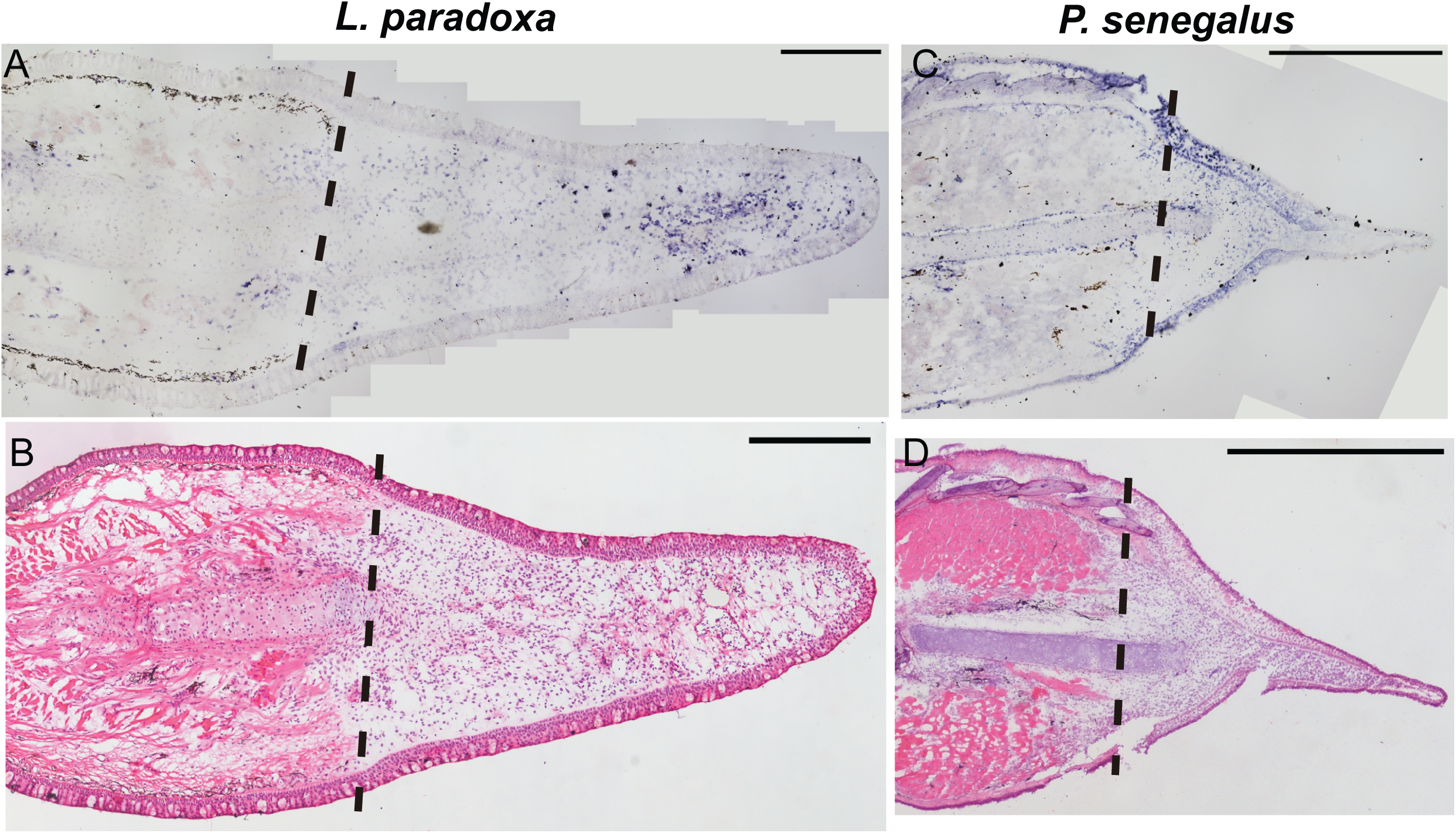
*vwde* is enriched in the regenerating fin of Lungfish (*L. paradoxa*) and *Polypterus* (*P. senegalus*). Expression pattern of *vwde* in the fin blastema tissues *of L. paradoxa* and *P. senegalus*. Longitudinal histological sections of fins from *L. paradoxa* at 21 dpa (A-B), and from *P. senegalus* at 5 dpa (C-D). (A and C) *In situ* hybridization using an anti-sense riboprobe to *vwde*. (B and D) H&E staining on sequential sections. All panels show posterior view, dorsal to the top. Dotted lines indicate amputation site (Scale bars, 1 mm in all panels).

To further investigate *vwde* during regeneration, we took advantage of the regeneration-competent and regeneration-refractory periods during *Xenopus laevis* tail development [16]. A blastema forms in response to amputation during both distinct developmental stages, but only in the regeneration-competent setting is full regeneration accomplished. This developmental feature provides an ideal situation to compare regeneration-component versus regeneration-refractory environments. We reasoned that finding factors that differentiate these two contexts may provide clues for identifying the core requirements for successful regeneration. We probed for the expression of *vwde* during the regeneration-competent and regeneration-refractory periods of *Xenopus laevis* tail regeneration. Interestingly, we found that *vwde* expression was present in tails prior to amputation in both the regeneration-competent and regeneration-refractory setting (Figure 3A-B, 3G-H). We found robust *vwde* expression along the peripheral edge of the amputation plane and near the blastema in regeneration-competent tails (Figure 3C-F). In contrast, in the regeneration-refractory setting, *vwde* expression was restricted to the peripheral edges of the amputation plane and was not detected near the blastema (Figure 3I-L). This indicated a striking correlation between *vwde* expression and regeneration, providing evidence that *vwde* may be an important factor in forming a pro-regenerative niche. These expression data across a range of species indicate that *vwde* fits the profile of an evolutionarily-conserved, regeneration-enriched gene and that *vwde* may play an important role in the blastema niche.

**Figure 3:**
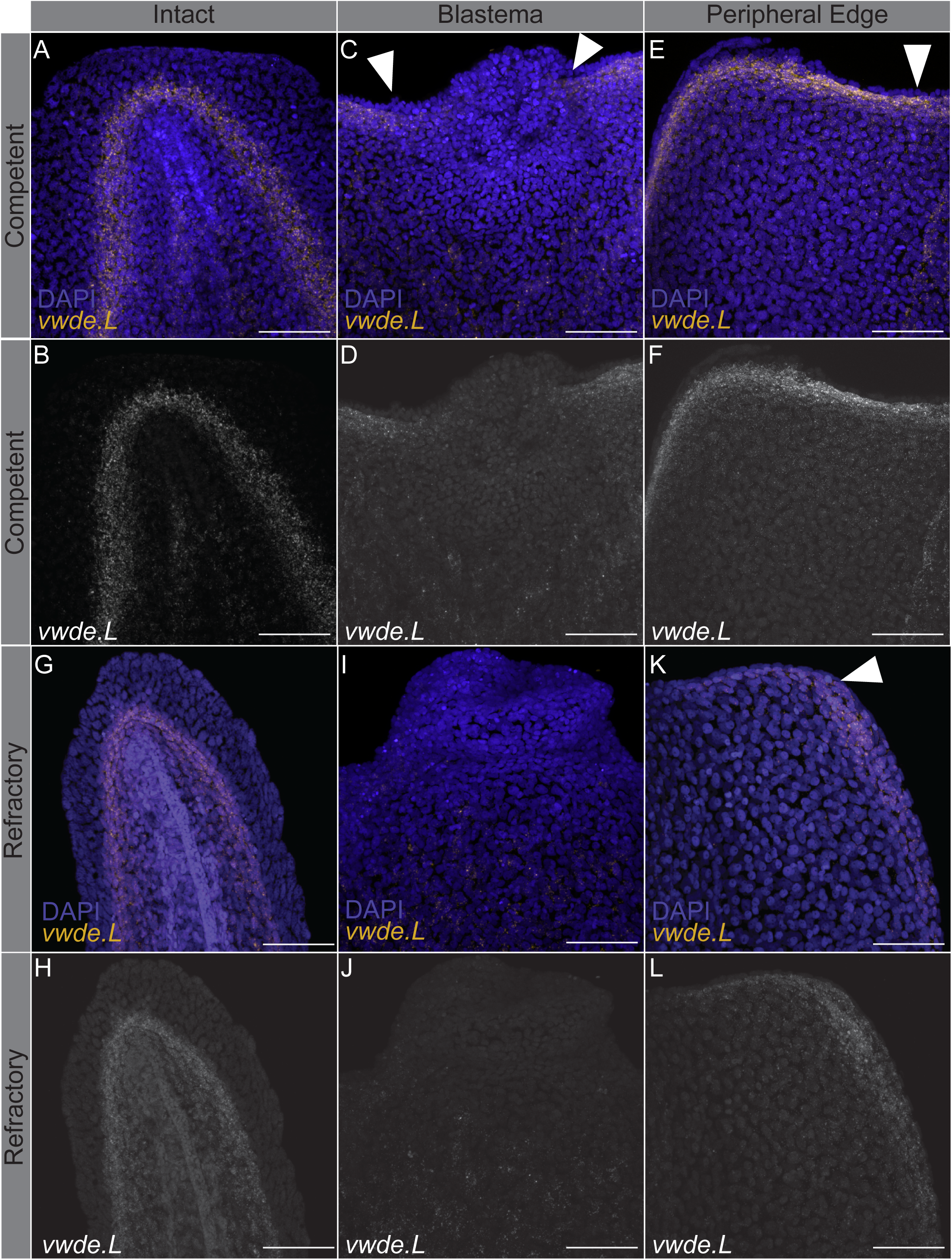
*vwde* expression is tightly linked with the regeneration-component environment. *In situ* hybridization chain reaction probing for *vwde* in (A-F) regeneration-competent *Xenopus laevis* tails (A-B) prior to amputation, (C-D) blastema 24 hours post-amputation, and (E-F) the peripheral edge of the amputation plane 24 hours post-amputation. (G-L) Regeneration-refractory tails (G-H) prior to amputation, (I-J) blastema 24 hours post-amputation, and (K-L) the peripheral edge of the amputation plane 24 hours post-amputation. White arrows indicate the location of *vwde* expression. Scale bars are 100 µm.

To investigate if *vwde* is required for regeneration, we performed morpholino-mediated knockdown at its peak expression in the medium-bud limb blastema. We found a substantial reduction in the length of the blastemas when Vwde was knocked down with two separate translation-blocking morpholinos (Figure 4A-B). Fluorescent reporter constructs with *vwde*-morpholino binding sites confirmed that both unique *vwde*-targeting morpholinos were capable of blocking translation (Supplemental Figure 3). Due to the dramatic reduction in blastema length, we investigated if *vwde* was important for blastema proliferation and/or cell survival. We found that knockdown of Vwde substantially reduced blastema cell cycle entry (Figure 4C-D) and did not alter cell survival compared to control limbs (Supplemental Figure 4). Due to the observed delay in blastema growth, we questioned whether blastemas treated with translation-blocking morpholinos were capable of recovering from the transient knockdown of Vwde and produce fully regenerated limbs. We therefore performed the same Vwde morpholino-mediated knockdown on a separate group of axolotls, and then allowed for the full course of regeneration to complete, harvesting limbs more than eight weeks post-amputation. We observed that one-time injection of Vwde-targeting morpholino caused substantial abnormalities in regenerated limbs, suggesting an essential role for *vwde* during limb regeneration (Figure 4E-G). We found defects in 4.2% (1/24) control limbs compared to 46% (13/28) of limbs treated with *vwde* MO1 and 25% (5/20) of limbs treated with *vwde* MO2 (Fisher’s exact test P < 0.05) (Figure 4F, Table 1, Supplemental Figure 5). A second experiment yielded similar results, with defects at endpoint in 27.6% (8/29) of control (*vwde* MO1 inverted) treated limbs compared to 47% (18/38) of limbs treated with *vwde* MO1 (Fisher’s exact test P < 0.05) (Figure 4G, Table 2, Supplemental Figure 6). We found a variety of defects, some of which are reminiscent of limb development phenotypes where limited distal elements are present such as has been observed in *fgf4,8*-double-knockout mice [17] and in the absence of *sonic hedgehog* (*ssh*) [18]. In addition, these phenotypes also resemble the defective regenerative spike characteristic of *Xenopus* limb regeneration [19]. Altogether, these data highlight the functional requirement for *vwde* during limb regeneration.

**Table 1.** Phenotypic outcomes of morpholino-mediated knockdown in axolotl (Standard control vs. *vwde* MO1, *vwde* MO2)

|  | Normal | Spike | Loss of distal elements | Oligodactyly | Polydactyly | Syndactyly | Additional elements |
| --- | --- | --- | --- | --- | --- | --- | --- |
| Control | 23 | 0 | 0 | 0 | 0 | 1 | 0 |
| *vwde* MO1 | 15 | 4 | 5 | 4 | 0 | 0 | 0 |
| *vwde* MO2 | 11 | 0 | 0 | 3 | 1 | 1 | 0 |

**Table 2.** Phenotypic outcomes of morpholino-mediated knockdown in axolotl (Inverted control vs. *vwde* MO1)

|  | Normal | Spike | Loss of distal elements | Oligodactyly | Polydactyly | Syndactyly | Additional elements |
| --- | --- | --- | --- | --- | --- | --- | --- |
| *Vwde* MO1 INV | 30 | 0 | 4 | 4 | 0 | 0 | 0 |
| *Vwde* MO1 | 20 | 6 | 9 | 1 | 0 | 1 | 1 |

**Figure 4:**
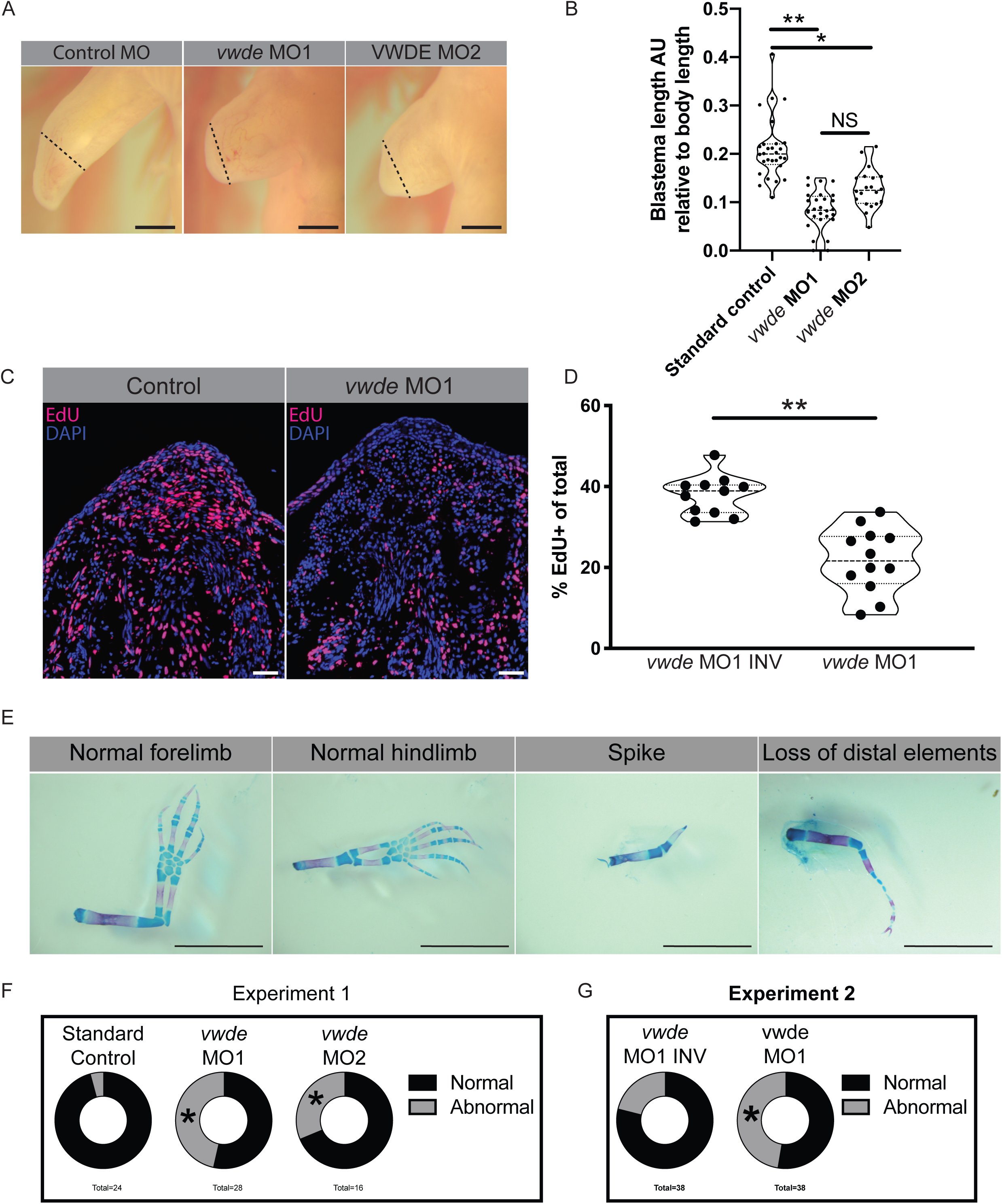
Vwde is essential for limb regeneration. (A) Representative images of blastemas 16 days post-amputation (9 days post-morpholino administration) from control morpholino (Standard control MO), *vwde*-targeting morpholino 1 (*vwde* MO1), and *vwde*-targeting morpholino 2 (*vwde* MO2). Dotted line indicates amputation plane, blastemas are all tissue distal to amputation plane. Scale bars at 1mm. (B) Quantification of blastema length 16 days-post amputation. Median and quartiles noted with dotted lines, ** indicates P< 0.01, * is P< 0.05 by nested T test. (C) Representative EdU stained sections of blastemas 10 dpa (3 days post-electroporation) of control morpholino (*vwde* morpholino 1 inverted, *vwde* MO1 INV) and *vwde*-targeting morpholino 1. Scale bars are 100µm (D) Quantification of percent of blastema cells positive for EdU in control (*vwde* MO1 INV) and knockdown (*vwde* MO1). Each dot represents a limb, 4-5 animals per group. Median and quartiles noted with dotted lines,** indicates P< 0.01 by nested T test. (E) Representative skeletal preparations of limbs after full regeneration after knockdown of Vwde at 7 dpa. From left to right, normal forelimb, normal hindlimb, spike, and loss of distal elements. Scale bars are 5 mm. (F) Donut plots of regenerative outcomes, pooled as abnormal versus normal from experiment with standard control morpholino, *vwde* MO1 and *vwde* MO2. Asterisk (*) indicates P< 0.05 by Fisher’s exact test comparing control versus *vwde* MO1 and control versus *vwde* MO2. (G) Donut plots of regenerative outcomes, pooled as abnormal versus normal from experiment with *vwde* MO1 INV and *vwde* MO1. Asterisk (*) indicates P< 0.05 by Fisher’s exact test comparing outcomes from *vwde* MO1 INV compared to *vwde* MO1.

## Discussion

Recent work, most notably next generation sequencing, has led to a plethora of information about the genes and cells that define the blastema [1,2,5–13]. However, it is difficult to determine which genes may have functional relevance based purely on their expression. We decided to investigate a single blastema-enriched gene, *vwde*, using an evolutionarily-informed approach, assuming that a gene whose expression is enriched in blastemas of multiple, distantly-related, species is likely a key factor during regeneration.

The *in vivo* assays used here place Vwde as a critical regulator of cell cycle entry during axolotl limb regeneration. Proliferation is a complex, but fundamental, aspect of regeneration, as there are many different cell types and potential origins of proliferative signals. Previous work indicates that mitogenic signals are produced directly following amputation independent of the nerve or wound epidermis [20,21], but are also provided by the nerve [22,23] or wound epidermis [24]. There are thus multiple sources of mitogenic signals in the regenerating limb, but it is unclear if mitogenic signals from multiple tissues are required simultaneously or perhaps in a more stepwise fashion to maintain blastema proliferation. Our data indicate that Vwde may be a blastema progenitor cell-derived mitogen, which adds to the potential sources of proliferative signals in the regenerating limb. It has been previously postulated that nerve-derived signals are required early on during blastema formation and growth, but a fibroblast-derived factor is required for complete regeneration [25]. We speculate that *vwde*, which appears to be expressed across the majority of cells in the blastema including likely fibroblasts, may provide one of the essential fibroblast-derived factors required after the nerve has provided sufficient input. While there is limited knowledge of fibroblast- or blastema cell-derived mitogens, *in vitro* cultures have shown that blastema protein extracts are able to drive blastema cell proliferation [26]. More generally, a global and temporally-based view of the cellular origins of mitogens and the cell types that require these mitogens will provide a better understanding of what is driving proliferation during different stages of regeneration.

In addition to the dramatic reduction in proliferation, we observed striking end point phenotypes after transient knockdown of Vwde. The loss of distal elements and spike-like phenotypes observed after Vwde knockdown suggest that Vwde plays a role in proximal-distal determination in the regenerating limb. These phenotypes showing similarities to *ssh* and *fgf4,8*-double-knockout mice, suggest that Vwde may be working similarly to—or in concert with—FGFs during regeneration. Though many FGFs are epidermal factors during limb development in mice and chick, FGFs are expressed in the mesenchyme during axolotl limb development [27] and regeneration [28]. Thus, it may be that during axolotl limb regeneration, blastema-derived factors are primarily responsible for proximal-distal patterning and that Vwde is working to promote the formation of distal elements. Intriguingly, *vwde* has remained unexplored in highly studied, but less regenerative species such as mouse and human, so whether *vwde* plays a role in limb development in these species is unknown.

It is interesting to speculate on what has been lost in amniotes that prevents appendage regeneration. One possibility is genes that are lost in amniotes and present in anamniotes can explain differences in regenerative capacity [29]. However, the absence of a gene in amniotes is not necessarily a prerequisite when considering which candidate genes might be responsible for high regenerative capacity. Alternative scenarios include, but are not limited to, genes that have lost ancestral pro-regenerative function or have altered expression domains/kinetics. *Vwde* may fit the paradigm of a gene that is present in both regeneration-competent and regeneration-incompetent species, but may exclusively be used in the blastema, a structure that cannot be produced by most regeneration-incompetent species.

While the blastema is required for regeneration, wound healing and activation of progenitor cells required for formation of the blastema must precede blastema formation. Based on the expression profile, we do not expect *vwde* to be a driver of blastema formation, but more likely a downstream effector once a blastema has been established. In most cases of amputation in less regenerative species, the blastema is not able to form, and thus we suspect that a more upstream or systemic factor may prevent blastema formation. While there may have initially been one primary cause of the loss of regenerative ability, such as the rise of adaptive immunity [30] or trade-offs associated with endothermy [31], it is likely that other aspects of the regenerative response have now been lost due to their lack of utility. If *vwde* played a relatively specialized function in the blastema and blastemas generally do not exist in less regenerative species then the use for *vwde* decreases. This could explain why 42.7% of human genomes have a predicted loss-of-function copy of *VWDE*, leading to speculation that *VWDE* is potentially drifting towards inactivation in the human population [32]. While the blastema remains the elusive feature required for appendage regeneration, this work illustrates that taking an evolutionarily-informed approach can lead to identification of functionally important genes. This also suggests that further work to understand the similarities between different species blastemas may help to elucidate the core molecular program of the blastema.

## Methods

### Animal Experimentation

All axolotl experiments were performed in accordance with Brigham and Women’s Hospital Institutional Animal Care and Use Committee in line with Animal Experimentation Protocol #04160. All animals were bred in house, but the colony was originally derived from animals obtained from Ambystoma Genetic Stock Center (Lexington, KY, NIH grant P40-OD019794). For amputations, animals were narcotized in 0.1% MS-222, confirmed to be fully narcotized by pinch test, amputated mid-zeugopod and the bone was trimmed. Animals were allowed to recover overnight in 0.5% sulfamerazine. For all functional experiments, all four limbs were amputated and injected individually. Functional experiments were performed on animals ranging from 3.8-8cm.

*Polypterus senegalus* and *Lepidosiren paradoxa* were maintained in individual tanks in a recirculating freshwater system. Animals were anesthetized before amputations: *P. senegalus* in 0.1% MS-222 (Sigma) and *L. paradoxa* in 0.1% clove oil diluted in the system water. Experiments and animal care were performed following animal care guidelines approved by the Animal Care Committee at the Universidade Federal do Para (protocol no. 037-2015). Pectoral fins in both species were bilaterally amputated. For *L. paradoxa* fins were amputated at approximately 1 cm distance from the body, and for *P. senegalus*, fins were amputated across the fin endoskeleton. Amputated fins (regenerating and uninjured) were used for histology, *in situ* hybridization and qRT-PCR analysis.

### Electroporation

Electroporation was performed while axolotls were narcotized in 0.1% tricaine and subsequently immersed in ice cold 1x PBS using a NepaGene Super Electroporator NEPA21 Type II electroporator. Settings for electroporation included: 3 poring pulses at 150 Volts with a pulse length of 5 milliseconds, a pulse interval of 10 milliseconds, a decay rate of 0 %, and a positive (+) polarity. Transfer pulse consisted of 5 pulses at 50 Volts with a pulse length of 50 milliseconds, a pulse interval of 950 milliseconds, a decay rate of 0 %, and a positive (+) polarity.

### qRT-PCR

Total RNA from regenerating or uninjured pectoral fins was extracted using TRIzol reagent (Thermo Fisher Scientific). Residual DNA removal and RNA cleanup were performed following the RNeasy Mini Kit (Qiagen) protocol. cDNA was synthesized from 0.5 μg RNA using the Superscript III First-Strand Synthesis Supermix (Thermo Fisher Scientific) with oligo-dT. For qPCR, amplification reactions (10 μl) prepared with the GoTaq Probe qPCR Master Mix (Promega) were run in a StepOnePlus Real-Time PCR System (Applied Biosystems). Gene-specific oligos (Table 3) for qRT-PCR assays were designed using Primer 3.0 (http://bioinfo.ut.ee/primer3/) and used in a final concentration of 200 nM to each primer. Each qPCR determination was performed with three biological and three technical replicates. Relative mRNA expressions were calculated with the 2^-ΔΔCT^ method [33], using *sdha* (*P. senegalus*) or *polrc1* (*L. paradoxa*) genes as endogenous control and the uninjured fin (mean ΔCT value of the three biological replicates) as reference sample.

**Table 3.**
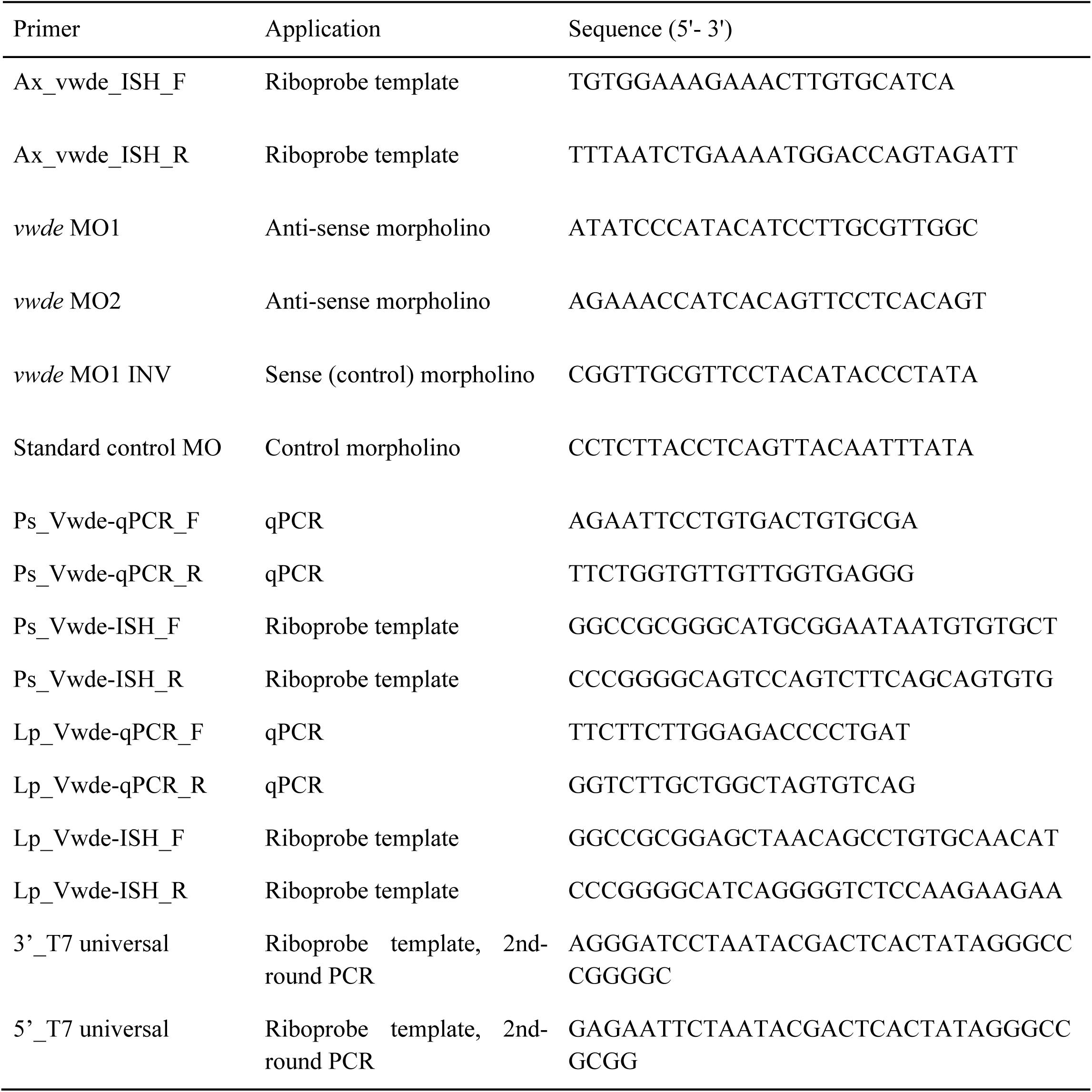
Oligos used for qPCR, morpholinos, and ISH probe templates.

### In Situ Hybridization

For *in situ* hybridization using axolotl samples a gene fragment from the 3’ UTR was amplified from blastema cDNA and cloned into the pGEM-T Easy vector and sequenced. Depending upon orientation, T7 or Sp6 polymerase was used to transcribe the probe. Primers for *in situ* probes against axolotl *vwde* (contig c1084387_g3_i1 from [11]) can be found in Table 3. Colorimetric in situ hybridization in axolotl tissue harvested from animals with snout to tail lengths of 9.5-11.5cm and was performed as previously described at protocols.io (https://www.protocols.io/view/rna-in-situ-hybridization-p33dqqn).

For *in situ* hybridizations with fish samples, fins of *P. senegalus* (5 dpa and uninjured) and *L. paradoxa* (21 dpa and uninjured) were amputated, embedded in TissueTek O.C.T compound (Fisher Scientific), and maintained at -80°C until use. Frozen sections of 20 μm were obtained on a Leica CM1850 UV cryostat, positioned on slides (Color Frost Plus/Thermo Fisher Scientific) and fixed as previously described [1]. Riboprobe templates containing a gene-specific segment (400-500 bp) and a T7 promoter sequence were produced by a 2-round PCR strategy (primers are listed in Table 3). Riboprobes were synthesized with T7 RNA polymerase (Roche) and DIG-labeling mix (Roche). Controls probes (sense riboprobes) were synthesized from a template containing the T7 promoter in a reverse orientation. A total of 300 ng of DIG-labeled riboprobe was used per slide during *in situ hybridization* performed as previously described [1]. Images were obtained on a Nikon Eclipse 80i microscope and processed using the NIS-Element D4.10.1 program.

### Whole mount RNA-FISH

*Xenopus laevis* eggs were obtained, fertilized, and cultured as embryos at 18 °C using standard methods as in [34]. All experimental procedures using *Xenopus laevis* were approved by the Institutional Animal Care and Use Committee (IACUC) and Tufts University Department of Laboratory Animal Medicine (DLAM) under protocol number M2017-53. Once embryos reached regeneration-competent (Stage 40) or regeneration-incompetent (Stage 46) stages, animals were anesthetized using 0.005% MS222 in 0.1X MMR and tails were amputated at the posterior third of the tail and allowed to regenerate for 24 hours. Embryos at both stages, which had not been amputated, were also collected as intact controls. Regenerating and intact control embryos were anesthetized in 0.005% MS222 and then fixed at 4°C, rocking overnight, in either 4% paraformaldehyde in 1X DEPC PBS or MEMPA buffer (0.1 M MOPS (pH 7.4), 2 mM EGTA, 1 mM MgSO4, 3.7% paraformaldehyde). We used a slightly modified whole-mount mouse protocol [35] using hybridization chain reaction v3.0 [36] with slight modifications. After overnight incubation, embryos were washed 3 times for 5 min in PBST and then taken through a methanol series on ice. This series consisted of 10 min washes on ice in ice cold 25%MeOH/75% PBST, 50%MeOH/50% PBST, 75%MeOH/25%PBST, 100%MeOH, and then finally stored in a fresh 100%MeOH solution. Dehydrated embryos were then stored at -20°C until use. For *in situ*, embryos were subsequently rehydrated via a reverse methanols series, on ice, with 10 min washes of 75% MeOH/25% PBST, 50% MeOH/50% PBST, 25% MeOH/75% PBST, 100% PBST, and another final wash in 100% PBST. Embryos were then digested with proteinase K (10µg/mL) in DEPC PBS for 5 minutes at room temperature. Post-fixation was then performed in 4% PFA in 1X DEPC PBS for 20 minutes at room temperature. Next, three five minutes washes with PBST at room temperature was followed by 5 minutes at 37°C in hybridization solution (50% formamide, 5x sodium chloride sodium citrate (SSC), 9 mM citric acid (pH 6.0), 0.1% Tween-20, 50µg/mL heparin, 1x Denhardt’s solution, and 20% dextran sulfate). Samples were pre-hybridized by full immersion in hybridization solution without probes for 30 min at 37°C. Hybridization was performed overnight at 37°C with samples immersed in hybridization solution containing twenty probe pairs against *vwde.L* (XM_018267342.1) diluted at 1:200 of 1 µM (hybridization chain reaction v3.0 RNA fluorescent in situ probes were ordered from Molecular Instruments (https://www.molecularinstruments.com/). The following day, samples were washed four times at 37°C in probe wash buffer (50% formamide, 5X SSC, 9 mM citric acid (pH 6.0), 0.1% Tween-20, and 50µg/mL heparin). Samples were then washed two times in 5X SSC at room temperature. Pre-amplification was then performed at room temperature for 30 minutes in amplification buffer (5X SSC, 0.1% Tween-20, 10% dextran sulfate). During pre-amplification, hairpin probes (ordered from https://www.molecularinstruments.com/) compatible with *vwde.L* probe pairs were heated individually at 95°C for 30 seconds and then snap cooled for 30 min at room temperature in the dark. After 30 minutes, probe pairs were added to amplification buffer at 1:50 (3 µM stock) and this probe containing buffer was subsequently added to samples, ensuring that samples were fully immersed. Incubation was performed overnight at room temperature. The next day, samples were washed for 5 min in 5X SSCT, twice for 30 min in 5X SSCT, and a 5 min wash in 5X SSCT. Samples were then stained with DAPI for 5 min in 1X PBS, washed for 5 min in 1X PBS, and then stored in 1X PBS. Samples were then mounted in low melt agarose and imaged on a Zeiss LSM 880 Upright. A median 3×3 filter followed by maximum projection was applied to all images.

### Morpholino design and administration

Morpholinos were designed and synthesized by GeneTools. Morpholino sequences can be found in Table 3. About 1.25 µl of morpholino was injected in the blastema and electroporation was performed as described in Electroporation. All morpholinos were 3’ fluorescein conjugated to allow for visualization. Morpholinos were reconstituted to 1 mM in 2X PBS and diluted to a working concentration of 500 μM in 1X PBS prior to injection.

### EdU staining

Stock solutions of 5-ethynyl-2’-deoxyuridine (EdU) dissolved in dimethyl sulfoxide were prepared per manufacturer’s instructions (Thermo Fisher). Axolotls (3-6cm tail to snout) were narcotized in 0.1% tricaine at 7 days post amputation and control or Vwde-targeting morpholino was injected and subsequently electroporated as described in Electroporation section of methods into the blastema. At 9 dpa, intraperitoneal injections with 400 µM EdU in 0.7X PBS at a volume of 20µL/g were performed. 18 hours later blastemas were harvested, fixed for 1-2h in 4% PFA and then taken through a sucrose gradient to 30% sucrose in 1x PBS. Tissue was then embedded in OCT and frozen in a dry ice/ethanol bath. Sections were cut at 16 µm with a cryostat, collected on Superfrost Plus slides (Fisher), and stored at -80°C. EdU staining was performed with the Click-iT EdU Alexa Fluor 594 Imaging Kit per manufacturers instructions (Thermo Fisher).

### TUNEL assay

TUNEL assays were performed as previously described [11]

### Skeletal preparations and scoring

Limbs were stained with Alcian blue/Alizarin red according to [37]. In brief, limbs were incubated with rocking overnight in 95% ethanol and then rocking overnight an acetone. Limbs were then incubated for at least 7 days in alcian blue/alizarin red at 37 °C. Limbs were then cleared by incubation in 1% (wt/vol) KOH, followed by 1% (vol/vol) KOH/25% glycerol, 1% KOH/50% glycerol, and 1% KOH/75% glycerol. Limbs were imaged in 1% KOH/75% glycerol. Alcian blue stock was 0.3% alcian blue in 70% ethanol; alizarin red stock was 0.1% alizarin red 95% ethanol; the working solution was 5% alcian blue stock/5% alizarin red stock/5% glacial acetic acid/volume in 70% ethanol.

Definitions for limbs after regeneration. Normal: All digits and carpals present, zeugopod and stylopod intact. Spike: Single outgrowth from amputation plane without obvious turn at joint. Loss of distal elements: Distal elements without obvious autopod. Oligodactyly: Loss or reduction in size at least one digit. Syndactyly: Fusion of digits. Additional elements: Extra bones in stylopod or zeugopod. For statistical analysis normal was compared to all of the above listed abnormalities.

### Ortholog analysis

The following proteomes were downloaded from uniprot.org, human (*Homo sapiens*, UP000005640, accessed 5/18/2019), zebrafish (*Danio rerio*, UP000000437, accessed 5/18/2019), mouse (*Mus musculus*, accessed 5/18/2019), amphioxus (*Branchiostoma floridae*, accessed 8/27/2019), chick (*Gallus gallus*, accessed 8/27/2019), sea squirt (*Ciona intestinalis*, accessed 8/27/2019), lamprey (*Petromyzon marinus*, accessed 8/27/2019), green anole (*Anolis carolinensis*, 8/27/2019), frog (*Xenopus laevis*, UP000186698, accessed 7/11/2019), frog (*Xenopus tropicalis*, UP000008143, accessed 7/11/2019). The South American lungfish transcriptome was downloaded from https://www.ncbi.nlm.nih.gov/Traces/wgs/?val=GEHZ01 and converted to a putative reference protein using TransDecoder (version 5.3.0) like so: ‘TransDecoder.LongOrfs –t’. The *Polpyterus* transcriptome can be found here: https://www.ncbi.nlm.nih.gov/bioproject/480698 and converted with TransDecoder as referenced above. The axolotl (*Ambystoma mexicanum*) predicted proteome was obtained from https://data.broadinstitute.org/Trinity/SalamanderWeb/Axolotl.Trinity.CellReports2017.transdecoder.pep.gz [11]. Cloning of axolotl *vwde* revealed a sequencing error in the axolotl transcriptome which eliminated the first ∼500bp of the sequence. We manually changed the axolotl proteome to include this corrected version of *vwde* (Supplementary File 1).

To predict orthologs, we used OrthoFinder2.0 (version 2.3.3) [14]. Orthofinder was implemented as follows:

‘orthofinder -f /path/to/proteomes -M msa -A mafft -T fasttree -t 20 -o /path/to/output/directory’

### Protein domain diagrams

The R package, drawProteins [38] was used to draw protein domains for different species Vwde. For all genes contained within Uniprot, these were downloaded directly with drawProteins. For genes not available via Uniprot (https://www.uniprot.org/)[39] (e.g. *Polpyterus*, axolotl, and Lungfish), the amino acid sequence of the protein was queried via Interpro with default settings (https://www.ebi.ac.uk/interpro/)[40] and positions and domain annotations were extracted and made into a matrix that matched the required structure for drawProteins. The Uniprot version of mouse *vwde* (Uniprot ID: Q6DFV8) in the proteome used did not co EGF-like domains, so we manually searched UCSC genome broswer to confirm this lack of EGF-like domains. This revealed a full length Vwde (ENSMUST00000203074.2), which was then fed into Interpro and domains were manually input into drawProteins.

### Vwde knockdown confirmation

Two separate constructs to test the target specificity of each MO used. GFP was removed and from pCAG-GFP (pCAG-GFP was a gift from Connie Cepko (Addgene plasmid # 11150; http://n2t.net/addgene:11150; RRID:Addgene_11150)[41] and replaced with vectors containing td-Tomato sequence and the morpholino binding site (Supplementary File 2). To confirm knockdown we co-injected and electroporated into medium-bud blastemas the generated constructs and the appropriate fluorescein-conjugated morpholino. Fluorescein fluorescence was used to confirm injection efficiency and td-tomato expression was used to measure ability to block translation.

### Statistics

Nested one way ANOVA was used to determine significance between blastema lengths. Each limb was considered a technical replicate within one biological (i.e. animal) replicate. Nested t tests were used to determine significance in EdU and TUNEL experiments, again treating each limb as a technical replicate and placing limbs from the same animal within one biological replicate. Fisher’s exact tests (control vs. treated) were used to determine significance of outgrowth phenotypes. Significant results were considered as P < 0.05.

## Contributions

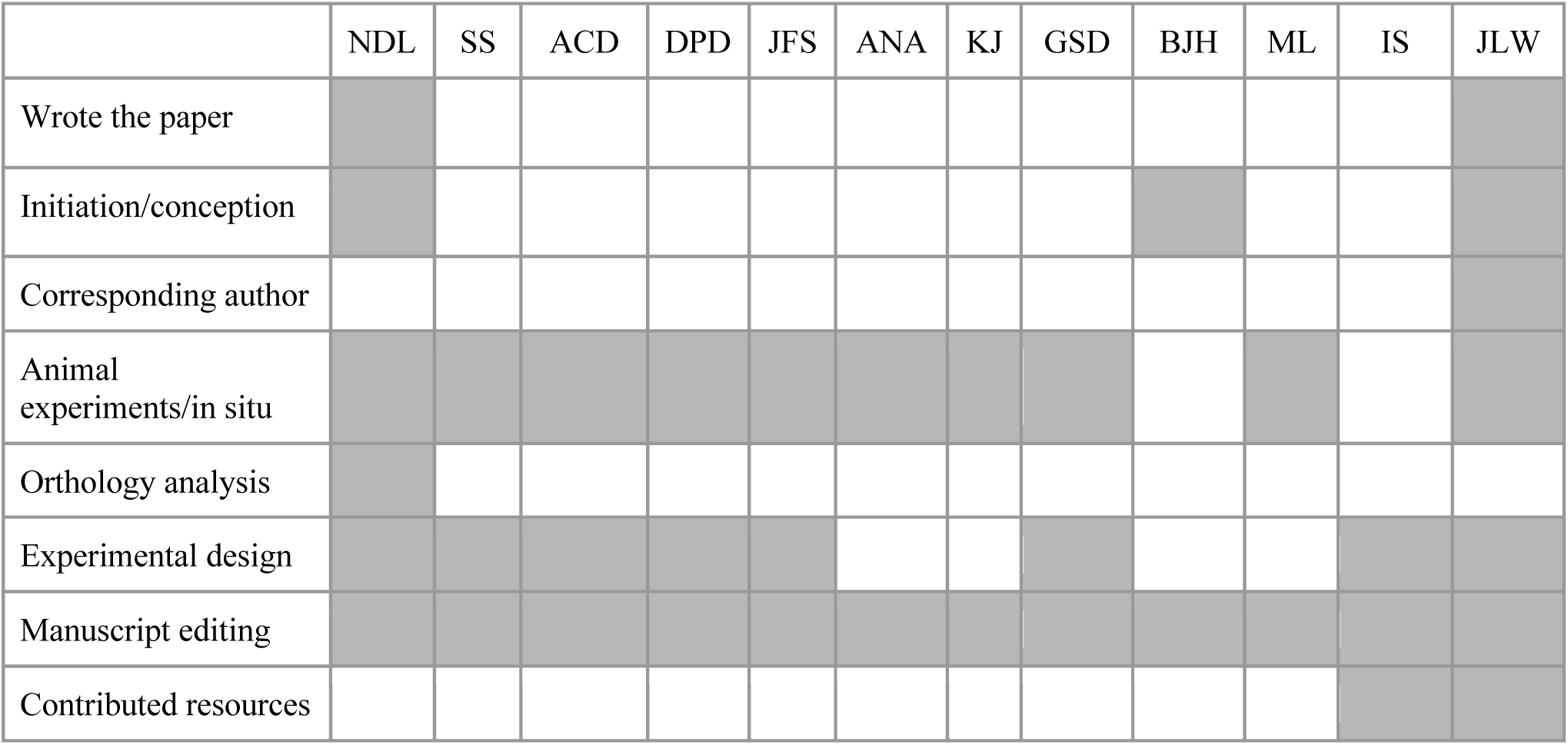

## Acknowledgements

This work was supported by the Eunice Kennedy Shriver National Institute of Child and Human Development of the National Institutes of Health (5R03HD083434-02 to J.L.W.). N.D.L. was supported by Award Number F32HD092120 from the Eunice Kennedy Shriver National Institute of Child and Human Development of the National Institutes of Health. Portions of this research were conducted on the O2 High Performance Compute Cluster, supported by the Research Computing Group, at Harvard Medical School. See http://rc.hms.harvard.edu for more information. Support for work on lungfish and *Polypterus* was provided by CNPq Universal Program Grant 403248/2016-7 and CAPES/DAAD PROBRAL Grant 88881.198758/2018-01 to I.S, and a postdoctoral fellowship from CNPq to A.C.D. We thank William Ye, Adam Gramy, Sarah Lemire, and Bonney Couper-Kiablick for expert animal care and other members of the Whited lab for feedback and discussions. We also thank Dr. Mansi Srivastava and Dr. Ryan Walker for their insights on orthology and protein structure and Dr. James Monaghan and Timothy Duerr for sharing information on whole mount in situ protocols.

