## Supplementary File 1 for "von Willebrand Factor D and EGF Domains is an evolutionarily conserved and required feature of blastemas capable of multi-tissue appendage regeneration"

Supplementary File 1
Sanger sequencing corrected axolotl *vwde* ORF
>c1084387_g3_i1_corrected
ATGTGCAACGGGAAGGGGAGAAGGGATAAGACCGAGGCGATGCAGGGCACCGAACCCTCCACCCG
GCCGAGGCTCTTACCGCCCCTGTGCACCCTGGTAAAGTCAGCCTTCGCACTCTGGATCCTTGTGAC
GTCTCTGGGGACAGCAGGCGCTCAGCAAGCTCCAGAGTGTAAAGCCAGCGGACATCAGATTCTGCA
GAATTCATACAGAAGCACTGATTTCGATTCCTTGAGACTTCAGCAGTCAGCCATTCAAGATCTGATCT
GCGACCACTCCCTGGCACCCGGCTGGTACCGCTTTGTGATCTTTGACAAGCCAGCAGAGATGCCAA
CCAAGTGCGTGGAGATGAACCACTGTGGAACCCAGGCCCCAGTGTGGCTTTCTCTGCGGGAGTCAG
AGTCCCTTCCTGGACCGGGAGAGATCAGGCAGCTCACGGCCTGTGCAACGTGGCAGTTTTTTTTCA
GCAGCTCGAAGGATTGCTGCCTCTTCCGTATTCCTGTGACTGTGAGGAACTGTGATGGTTTCTATGT
CTACCTACTGCAGCCAACGCAAGGATGTATGGGATATTGTGCAGAAGTTGTTTCTGATACCACATTG
CCAACATGTGGCCCTGATGAAGTGGATATTGGTGGTGTGTGCACTGGTAAGCAGTCATCCCCCCTGT
CATTACCACCATCAGCAATGCTCCCACCGCCACCAGGTGTCCCAGAAATTGTGGCAGAGTTGTCCG
GGTCTAGCATTTACCTTAAGTGTTTCTTTGAAACCTCTTCCACAAACAGCTCGGTGGGTTTCGTTGTG
ACCTGGTCCAGGCTTTCTCTTGATGAGATCAAAGAGGAGCTTCGGCAGGAGACCACGGTTCAGGCG
TTCTCCCTAATAGAGCTGGATGGCATAAACCTCAGGCTCGGAGATCAGATTTATTGCAGCTGTTCTA
GTTTTTTTCTGGAACAACCAGAAGTGCAAAGCCAGAGCATGGAGAGCAGTGAATTCTTTGCTGGTGT
CAAGTTGCTTCCTGAAACCTTTAGTATTTCTGAAGATGGACGAGAGCACCTACTGACCGTTGAAAGTA
CGGTACCGATTCCTTGCTCAGAAGTCAGCCAGTTGGACCATGAATGCAAAGTGTTGCTAAAACTAAG
AACAGTCAACGAAGATGAAGAGCAGCCTGACCTTAATGTGGTTCTTTCAACTTGTCGAGTAGCTCTG
CGGCAAATGTATTGCCTTAACGGTACATGCAGTCATGCTAATCTACTCTTCACTGCAGTGACAGACTT
TGCCCGAGATGGAGACAGAGCTACACACATTGCTATTGATCCAATCATTAGTGACAACTTTCTCTGGA
ATGGATATGCTCCAAAGGGAATACAGATTACAGTGAAGGATTTGCCAACTGCATTTTGCTACTCATTC
ACAGACCCTCATATAATTACATTTGATGGCAGGCTTTATGATAATTTTAAAACAGGAACTTTTATGTTA
TACAAAAGTGCATCTCGAGACTTTGAAGTCCACGTGCGGCAGTGGGACTGTGGGAGTCTCCACTTTC
CGGCATCATGTAACTGCGGCTTTGTTGCCAAGGAAGGAAGTGACATCATTGCATTTGATATGTGCAG
TGGTCAGCTACATGAATCACAGCCCCACTTGTCTGTAAAGAGCAAGGACACTACCAGTGATAATGTC
AGAATCACCGAATCCTACTTGGGAAGAAAAGTTACAATTGCCTTCTCCTCTGGAGCTTTTATCCGTTC
TGATGTAAGTGAATGGGGAATGAGTCTGACCCTTCGTGCACCAAGCTCAGACTACAAAAATACTCTG
GGACTCTGCGGCACTTTTGATGGAGAGATGGAGAATGATTTTCATGACAGTGATGGGTTTGAAATCA
CACAGACCACCGATAATCAACACACGTTTATAAATGAATGGAGAATTCCTCCAGGGGAGAGCTTTTTT
GACAAAACTCCCACATCTTTGACTTCGCCCAAAAGGAAGCATTTTTGCGGATGTGCCGTTGATGACG
CTGAGCCGTATCAGTCTTCAAACCGGCCGGGGCCTGGATCCCAAGATGACTTTCTGGCCCCGTGTA
AAGGAAACAGCAATGTCCAGCAGTCTACCCTGATACCGGGACTAGATGTCACTGCAGAATACATCAG
CTCTGTTGAACACAGCCGGGATTTGTCCAGACGGTCACCGCCCGATGAAACTGATTTGGTCAGCTTC
ATTTCTGGAAATAAACTGCAATTGAAAGAAACAAAACAAGTAACTGCTACAACACAACAGATGGCCAC
TCCTGCAGTGGATGAAGACGACAGAAAGGGCGAGCATGGAAGCCATTCCTCTAAAGGACAGTCAAT
CAAAAATAGATTAAAACGTCAGAATTATTATGAATACCTGCCCACATTTCCGTTTCAAAGCCTGAGTCA
AACTGACCTTGAAGGATTCAACTATTTTTTCCCAGAGGACCACACGACGGATACGCAGCAGGAGTAT
ATGCCCTCCTGGCCTACACCGTCTGGTCTTACGCATGCCGGTGCATTGGAGGCATGCCAGCAGACC
ATCAGCAACTCTAGCATCGCCAAATCTTGTGTGGACCTTCTTGGGGGCAGGATCGGGGATGTTTTGG
AGATGTGTGTTACAGATTTGTTGCTGAAGGATGACCTCACCTGGGCAGAGGCAGGCCTCCCGTTGC
TGGAGAATGATTGTGAGCGCAGGATTTTGGAAGAAGGGAGCTACAATGCTGAAAGATATGGTGATGC
ATTGGAAAGTATGCTCCAAGCATTAAAGTGTCCCAAACTCTGCAGTGGAAATGGACAGTGCTTGGAA
TGGGGATGTGCCTGCTTTGAAGGATTTAGTTCTTATGATTGCAGTATTCTTTCAGATCAGATCCCAGA
AATTATAGAACTGGAGAATGCTGGTTTGTGCGATGTTAGGCAATATGACTGCACATCTGTCAGAGTGT
ATGGTCAGGGATTCAGGGAGTCACCCAATCTGAAATGTGAAGTTGCAAAACATCAGTACTCCGATGG
TAAGTGGTTTCTGGCTGAGCCTGAGGTTATGCATGCTGCGTTCCGGAACAGCAGGGCCATTGATTG
CCAACTGCCGTCTGATGGGCAACACTCGGATGGTGTGGATCTGGTGGATGATAAACCGATTGTGAT
GTGGCAAATCAAGGTTTCTAATGATGGTTATGTCTACAGTAACCCTAAAATAATGTCGCTGTTTGATG
GAGCCTGCCAAACCTGTGACCCACAATCTGAAGGTTTATGTACTTTAAAGGAGAAAACCTGCAACAT
CGAAGGACTCTGCTATGGAGAAGGAGACCCAAACCCCAGCAGCCCCTGCTTACTTTGCAGACCCGA
AATATCCAAATTAACGTGGTCCATCTCTGAGAGCAATACAGCCCCTGTGTTTCAAAAGTTGCAACACA
AGCTGCAGACATTTTATGGGGAAAACTTTGTGTATCAGTTTATGGCGTCCGATCCAGAGGGCTCGGC
CATCTTGTTCACAATGGACAAAGGACCTGAAGGCTCCAGTCTATCGCCTGCTGGACTTCTCATTTGG

AAAGCCATGTCAAAGGAGCCACAGATGTTTCATTTTTCTATAACTGATGACTGCAACGCTGAAACTAA
AGTTTCTGTAGAGGTCAGTGTGAAAAGCTGCAATTGCATGAATGGTGCGTCTTGTGTAACAAATATTA
ACTTCCCTCCTGGGAGAGGAGAATACCTATGCTCATGTTTGGCGGGATTTGAGGGTGAAGATTGTCA
AATAAACATTGATGACTGTACATCCAACCCATGTGGCCTAGGCACATGTTTCGATGAAATAACCAACT
ACAGATGTGAATGCACAGCTGGACTCACAGGTAGAAATTGCCAAGTAGATGTTGATGAGTGCGCGTC
CAGCCCTTGTTTTCCTGGAGTCATTTGCTTGAACACATTTGGTTCCTATCATTGTGGCCCGTGCCCCA
GTGGGCTGGAAGGGGATGGAAGGGTCTGCTTCGTGGAAACACGACTACCAACAGATACCCCTTCAC
ATTCAGTGCTCGCAGAAGACTTAGAATCAGATTATGGTGACGTTGAAGCTGATGCAGAGGATAGTGA
AACAACTGAATATTATCATTCAATTCCTGAAGAAAATGAAACCACCGATGAACAAACATCTGAGTCTG
TTTCTGTGCAAAGCAACATTTTGGCAGGTTTGAAAAATCCATGTGCTGCTAATCCCTGCTTCCCTGGA
GTTCAGTGCTTTGAAGACCTATATTTAGGAGACCATTACACATGTGGAGACTGCCCACCAGGCTTTTC
TGGAGATGGCCATATATGCAGAGCTCATGTGGCATCACCTGAATTTCAGCCTCTCCCGATAGACCAT
CCAGGAACGGGCATTGATAGAAGTGTCACTCCCAGTCCAGAGGCTGCAAGGAATCTGCTGTCACCT
TCTCCCCCTGAAGCACAACCGCCATCTACCACATCAAAGCCACTTGATGTCCAGGAATCCCTTAACA
GCCGTTCTTCTGCAACCAGATTCATAGGGCGGGGCTCTAGGACACATGAGAATTTGGACCCGTTGG
GCACATCTTTTGTAGGGGAGACATTGGCTGGTACCAGTGGGATCAAGGATGTATTTTTACCAGAAAA
AGACAACAACATATGGACTCAATTTAAAAATCTGACAACGACCCACAAAGAAACTCAATCCCATGCTG
AAACAAAACTGAGCGAAGTGAGCACTTTAAAAGAAAAGCTACCTGTGAGAAACACTGGTTCCAGATTT
CAGAACCCAACAGTCAGGCAAAGAATTGGCTCTTCACATCCAATTTCTGTAAAGATCACAGATGCTCC
CGAAGAAGGTGAAACACCAGTGACAGCGGTCTACAAGACAGTGACGTGTGCAGACTTCCCATGCTT
TCCGGGTGTGCCTTGTGAACCCAGTCAAGAGAGAGGATTCAAATGTGGCCGCTGTCCATATGGGTA
CCATGGCAACGGCGCGACTTGCACAGCAATATGTAGGCAGCCATGTGGTAAAAACATGGAGTGTGG
TGCGCCAAATACATGCCGTTGCAAGCCTGGTTATTCTGGATACAACTGCCAAACTGCTGTGTGTCGA
CCTGATTGCAAGAATCGTGGAAAGTGCATTAAGCCTAATATCTGTGAATGTGCCCCAGGATATGGCG
GTCCTACGTGTGATGATGCATTTTGTAATCCACCTTGTCAACATGGGGGTGCTTGCCTGGCTCGAAA
TGTTTGCACCTGTCCCTTTGGTTACGTCGGACCCCGATGTGAAACAATGGTTTGTAATCGACACTGT
GAAAATGGTGGCCATTGTGTTGCTCCAGAGGTCTGCAAGTGTAAATCTGGCTGGTATGGACCAACAT
GCAGCACAGCACTTTGTAGCCCAGTCTGTCTCAACGGTGGAACGTGTATAAAGTCAAATGTTTGCCT
TTGTCCGAATGGATTCTTCGGTACTCAATGTCAGAACGCGGTCTGTAACCCTCCTTGCAAGAACGGT
GGCCACTGCATGCGGAACAATGTCTGCTCCTGCCCTGATGGCTATAATGGCAAAAGATGCCAAAAAA
GTGTCTGCGATCCAATGTGCATGAATGGAGGACGGTGCGTAGGGCCCAACCTGTGTTCATGTCCCT
CAGGGTGGAAAGGAAAAAGATGCAACACCCCTGTCTGTCTTCAAAAATGTAAGAATGGTGGGGAATG
CATAGGACCAAACCAATGCCATTGTCTTCCAGACTGGGAGGGAATGCAGTGCCAAACACCTTTGTGC
AACTTAAAATGTCTATATGGAGGAAAATGTGTGTCGCCGAATGTCTGCTCATGCCGTCCTGGATACA
CTGGAATAGGTTGCAGTAAGAAGCTGCAGGTACAAAGTCGTCATGGTTGA
>PS64836_c0_g1_i1
CTTGGAAATACAGCTTAATAGAATAACAAAAAATAAACAACAAAACGTTTATTCAAAAGA
AATAATGACATATTCACATTATTTTACAAAATTAAAGTAACACTAAAGGCACAAAATGCA
TTCAGTAAGTAACAAATAAGAGTTGAGTGCAGTAAGGCACAATTTTATTATAACCCCAGA
GGAGCCACACAATGAGAATCTTAAAGCTGTTATATTTAGTTATACTTCGGGAAAGACTGT
AATCAAAGTTCATTTAAATTTGTGACATTTTTTAAATAATGGATGTCTACATTTTAAACC
TCATAAACCTACAAGTCAAAATAGGCGTGATATAAACTAAAATAAAACAGTATTTAAATA
GAAATCGTCTATTTACCATGTACACGATTTACAAGTTAAGAAACACATTTCAAAGAATAA
CTGAAATTGTCATATTTAGCATTATTATACATATCTAAAAACACAATATACCATTTTTGG
TTTCCTTGTCAACAAAATACTCTACTTTTTCTATTCTGTTTCTGTGCAAGCTTAACTTTT
GCCCATATAGCGTAAATCAATCGAGATTTGCAATTTTGTTAAAATATAAACATATTTTGA
CGTATTATAGGTAGGCTCCCTTTATGTTTTGGGTTTAATTGATTTCAAAAGTAAAGCTAT
TTTTTATGATTATGGATGTGATTTACTGTAATATATTTATACTACTATACATATCCATTT
CAGAAGATAAAAATTATGTCAATTTGAAATTTTATTAGATCATAAAATAAATTTGGAAGT
GGTCAGTTACATATAATAATGTACACCAAGGCCATAGTGTTTTAAAAAGCAAATGGGCAT
TATGTTTGATGTCTACTTTCACAGTTAAAATTAAACAGAATGATTAACTCGTCCACTCAT
TTTTCAATAACCCGCAGCAAATTGTTAAATACATATATACATATACATGTGAAGTAGGGG
ATGAATAATTTGAAAAGAAAATGAAAGTCTGAATTTAAAAATAGAAATTTTTAATTAACA
GAATTTTCTATGCATGAAAAGACTATATGTGGATTTTAATTTGGATGTCATTTGAATATT
TCCTTCAAATTGTCTTCTTCTGAGTGAAGTCCAGTCTTCAGCAGTGTGTGTTAGTTAAAT
CTGTATCAGCAATAATTAGCTTTTTGTTACTTCAAATGGTACTGCACCTTTGTACTACAG

AAAGCCCCAGAGAATCCTGGACGACAAAAACATCTGTTAGGATACACACATCTGCCCCCA
AACAAACATTTATTGTGACATATTGGTGTTTGGCACTGTAGCCCTTCCCATCCAGCTGGA
CAGTGACATGTATTGGGTCCCACACATTCTCCACCATTTTTACACTTCTGTAAACAAATG
GGAATGTTGCATCGTTTCCCTTTCCATCCTGAGGAGCAGGAGCAAACATTGGGTCCTACA
CACTTTCCACCATTCATGCACACTGGATCGCAGACACTCTTTTGACACCTCTTTCCACTG
TATCCATCTAGGCATGAGCACACATTATTCCGCATGCAATGTCCACCATTTTTACAAGGT
GGGTTACAAACAGCTAAAGGAAGCAGAGTTTTATGTCATTAAAGAACATTGTACTGCATC
AAAGTTACATCTATTTAAAAAAGCATATACACTCTTACCGATCTGACACTGAGCTCCATA
AAAACCAGCAGGGCACATGCAGACATTTGGCTTAACACATACTCCACCATTTAAACATAC
AGGTGTGCAAACAGCTGAGCTGCAGGTTGGTCCATCCCATCCTTCCTTACATTTACAGAC
ATCTGGGCTAACACATTCTCCTCCATTGTCACAATGTCGGTTACACACCATTGTTTCGCA
TCTGGGACCAACATAACCATAAGGACATGTACAAAGGTTTCTTGCTAAGCATGTACCGCC
ATGGAGACAAGGAGGGTCACAATTTGCTTCCTCGCAAGTTAGCCCCCCATATCCTGGAGC
ACACTGACATACATTTGGCTTTATGCATTTCCCACGATTTTTACAGTCTGGACGACAAAC
AGCGATGTGACAGTTATATCCAGTATACCCAGGTTTGCACCTGCACGTGTTTGGTAAAGA
ACATTCCATATTCTTGCCACATGGATATCTGCAAATTGCTTTACACTTGGCACCATTTCC
AGAATAACCATATGGACAGCGGCCGCATTTAAATGACCCATGTTCTGATGGTTCACAAGG
GACACCAGGGTAACATGGAGAATCAGCACAGGTGAGCAATTTTGGAAATGATGTAGTCCT
TTCTTGTGATTCATATAAGTCTTGTGGAAGAACAATGGCAGAATGCACCTTAGATCCTTT
ATATGTTATTCCAGAAGTTCTAGTTTTTGGAATTCTTTTGTCTTCATTCACTTCTTCCCT
AGAAATGTTTCTTTTATTACCAATGTTAAGCAATGTATTCCTACTATTTAAATATGGCAG
GCTTAGTTTGGGTTTTGCTTGAGGAGACTCAGGGCGAGAATTCAAAGTCCTATGGAAAGA
CGTTAAAAGTGGGATTTCCCCAGTGGTCTTTGTTGAGACTGTAAACTTTCCAGCTTTTTT
AGTTGTGGCTGAATGCTTGGATATTGAATCTGTAGTCATCGTAATTGTGGGTGGAGTAGG
TAAGTTTACTACATTTAGTGTAGCTGTATCAGTTGTTGTACTTCTTGATCTTTGTCCTGT
CAGAGTTAACCTGCTGCATGTAAAACCATTACCATGAAATCCTGGAGGACAGCCACCACA
AGAAAATCCTGATTCTTGTTCTGGATCTTCAAAGCATTGCACACCTGGGAAACACGGCTT
AGCAGCACAACCCTTTTGAGTGCCAGAGCTAAATTTACGTCCTGGTGATTGCTGACGGTG
AACACTGGTTTGTTTCTTTGTTTCAGTTATTTCATTTTCAGCTGGGGTACTGATTTGTGT
TTGTATTTCTGGCCCAGATTTATTGTCATTGCCTTCCTCCTCCTCAGGATACACCTTCTT
AACCTCTTCCTCTTTGACAAAAGGATTCAGATTCAAAGATTCTTCGACCGCTTTACAGTG
TCTGCCATCTCCTCTGTATCCTTCTGGACATGAATCACAGATGTAGGATCCAAAAGTGTT
ATTACAGCGAACTCCCACAAAGCAGGGCTTTGATGAACATTCATCAACATCCTCTTGACA
TTGTTCACCTGTAAAGCCTGAGTAACATTCACAAGAATATGCATTGAGGCCGTCCACACA
CCTGCCGAAGATGCAGGGATTTGATTTGCAGTTGTCAGAGTTGAAGTTACAGTAGTCACC
TTCAAATCCTCGGGGACAGACACAAACATATTTTCCACTTCCAGGAGGAAAGTTAATATT
TGTTACACATGTTCCTCCATTTAAACACTTACATGGCTCAACAATTACCTCTACAGATGC
TTTAGCTTCAGCATTGCAGTCATCTGTTACAGAAAATCTGAATGTTTCCGTTGTTTCAGA
TGTCACCTTCCAGATCAAAAGTCCAGCAGGTGAGAGTACTGCCTCCTTTGGACCTTGGTC
CAGTGTAAAAAGAACTGCAGAACCTTCAGGGTCTGTGGCCAAGAACTGGTACACAAAATT
TTCACCTCTGAATGTTTTAAGCTTTGCTTGTGATGGTTGGAAAATGGGTGGCTCATTTTT
TTCACTAATTGACCACGTAAATTTTGAAATGTCTGGTTTACACAGCAAACATGGACTTAT
TGGATTTGGGTCACCTTCTCCATAGCAAAGTCCATCAATGTTGCAAGTTCTTTCCTTTAA
TGAGCACAGTCCGTCAGAACTGGCTTCACATATTTGACAAGCTCCATCAAACAGTGTTAA
TGTTTTTGAATTGCTGTATTCATAACCATCATTTGAGACCTTAATCTGCCATCTGGCAAG
AGGTTTATCATCAACTATATCAATGTCCATATTATGAGGTGAATGATCATTTTCCCTTGG
TAACTGACAGTCTAAAACAGTGTTGCTCAAGAAGGAAGCTGTTGTAAGCTGGGGATCACC
TAAACTCCATTCATCAGAAATAAACTGTTCTTTTATGACTTCACACTTCAAGTCTGGTGA
ATCTTTAAAACCTTGACCAAAAATTCTCACTGACGAGCAGTCATATTGACGAACATCACA
AAATCCAGCATTTTCTAGCTCAATAATTTCTGGAGGCTGATCAGAAACTATACTGCAGTC
ATAGGCCCCATATCCTGGATAGCACACACATCCCCATTCGGCACATTCTCCATTCCCATT
ACAAAGGTTTGGACATTTCAGTAACGACAAAAGGCTGTTATACTCCTTTGAGTGGCTCTG
CTCTATAAGCTTTCTTTCACATTCATTCTCTAGAAGGGGCAATCCTGCTGTTGACCAGCT
GATGTCATCTTTTAGTTGAATATCTAGAATGCACATGTCTATAACATCCATTACTCTTTT
ACCAAGAAGGACTCCACATCCCTTGCCTATGCTTGAATTGGCTACTGTATGTTGGCACAA
CTCAGCTACTTTTGATTCTGTCAGTCCTGAAGGAGTTGGCCAAGAAGGCACAGTGTCAAA

GGCAGAATTAGGGGTGTGGTCCTCAGGGAAGAAATAACTGAAGCCTTCGAGGTCTGTCTG
GCTGAGACTTTGATAGGGAAAGCTTGAAACATATTCATAATAATTCTGCCTTCTTCGTCG
ATTGTTAGGCTGCCACTTATTCTGCAGATCATCATGTTTATATGCATCAGTGATAGCACT
TCTGCTGGGTGAACCTCTTTCCATTGTAATGTATTTATGTGGCCTAGCGAACAGATTACT
GTGTCCTGATGATGTTTCTTCTGAATTGCTTTGATTATTTTTATGAAGTACTCTTGAAAG
GGGAGAAGAAAAGGAATCTGATCCCGTCCCAATCTCTCGTCGATGTAATTCCCTGAAATG
CTCCGTTGGATTTATGTATTCGGCAGTTATGTCCAAAACTGGAATTAAAGTTGAAAGTCT
TACATTTGCATTGTTTAAACAGACTGGTGAAGCCCCTGCATGTGCGGGAAAATTAAACTT
GTTTGATGGAGAATGTGTCTTCAGTGTGTCAGTTTTACAGTTACAATAGTGCTTTTTCAG
GAATGTACTTTTAAATGATGGTGAATGATCAAATAAACTATTTCCCGCAGCCACCCTCCA
TTCCTCAATAAAAACAGATGAATTTTGAAGTACAGAATGGCCATCTGCACCGCGGAAGTC
ATTTTCTATATTTTGATCAAATGTTCCACACAAGCCTTCAGTATGATGATGATCCAAGCT
AGGAGCTCGGACAGTTAAACTCATCCCCCACTCATTTACATCAGCTCGAACAAAAGCTCC
TGATGAAAATGTAACCGTGACTTTTCTTCCTTGGTATGATTCTGTTATGATGATATTTTT
ACTACTAGAGTCTTTGTTCTTTATTGCTAAGTGTGGTTTTGTCTCACGTAGTTGACCATT
ACACATGTCAAACGAAATCATATCTTCATCTTCTTTAGCAACAAAGCCACAGATGCATGA
AGCGAAGTAACTGTGGCTTCCACAGTCCCACTGGCGAACATGCACTTCAAACTCCCGGCC
CGTACTTTTATATAATACAAATGTTCCAGTTTTGTAATTATCATATTTTCTCCCATCAAA
TGTAATAATATGAGGGTCTGTGAATAAATAGCAGTATGCAAAAGGTGCATCTTTAACTGT
AATCTTGGTGCCTTGTGGAGAGTAGCCATTCCATAAGAAGTTATCACTAACAATGGGTAG
GACCTCAATTTCTGATATTTTGTCAGAATCCATAATAAAATCAGTAACTCCTGTGAAATA
TGTAGTCGCAGAGGCACATATTCCTTTATAACATGGAGTATGTTGTAATTCCACTTGACA
AGAAGAGAGAACAAGATCTGGTCTCAGATTGTCTTCATTTAGTGTGTGCAGTTGTAAAGT
GATCTTGCAGTCCTCTTTAGGTTGTCCAGATGAAGAACAAGGAATCGGTACTGTACTCTC
AATGTGTAATTTATGTTCTTTACCATCTTCTGATATAGACGCAGTTTCTGGATGAAGCTT
AATGCCAGCAAAAAATTCTTTGCTTTCAGCTGGAGGTCCTTGAATATCAGGAGAATCCAA
GAAAAAGCTGGATATTTTGCAGTAGATCTTGTCTCCAAGTCGAAGATTTATGCCATCTAG
CTCAATGAAAGAAAAGCTTTGTACTGTTGTCTCTTGCTTCAGTTCTTCTTTAGTTCCATC
AGAAGAGAGTCGAGTCCAAGACACAACATGTCCAAGAGAACTATTTGTGACAGGACTATC
AAAAGAGCATTTTAAAGTAATGGTGCTGCCAGTCAGCTCAGCCACTACTTCTGGTGTTGT
TGGTGAGGGTGGATATTTAGCTTTACAAGTTCCACCAACATCTGTTTCAGCCGGGCCACA
GAGGTCTTGCTTTATGTCTGAAACAACTTGAGCACAGTAGCCCATACATCCCTGTGTTGG
CTGCAGTAGATACACATTAAATTCTCCACAGTTTCGCACAGTCACAGGAATTCTAAACAG
GCAGCAGTCCTTTGTAGTGCTAAAGAAAAACTGCCATGTTGCACAGGCTGTGAGCTGCTT
CACCTCCCCAGGCCGAGGAAGAGACTCTGATTCTTTAAGTGAAAGCCACACAGGGGCCTG
TGTGCCACAGTGATTCATCTCCACACATTTTGTTGGCATATCGGCCGGTTTGTCAAAAAT
ATGGAAACGATACCATCCAGGAGTTAAAGAATGATCACATATTAAGTCCTGTATTGCTGA
CTGCTGAAGCTTTTGAGAATCAAATGTAACACTTCTATAAGGGTTTTGGAGAATACGATG
GCCATTGGGGCTGCATTCAGGAGCTATTTGACAGTGTCCAGCGGTCATCGCCAGAACGAT
CATGGTAATCCTGATTGCAACATATTGTACCAAATGTTCTAGTGCCATGGTGAGTGACAT
CCGACGTGTCTTTAGTCTCCGGTCCCTCTCACTTAACCCGATTACTCTGTCTGCTCAGAT
CAGCCAGTATTCCAGTGTTGTTTCGTGAGAAGTACAGACAGCAGCGCCGCGTCTCTCACT
GAGAGCAGTAACTGCTCGAGCGCAGCTGGACGCTCCACTTTGTCGCTCTCACACAGCTGG
CGATGCGGGTCGCTTGAGCTCCCCAGTGAGCTGAGCCGGCCGTGTGAGATCGCGTACCGC
GCCTCAGTAACAGCTAAGGGAGGAAAATCTTCGCAGCCTAATTTGGGCAGTTTAATGCAT
CAGTTCTCGACTCACTGGGCGATGCGCACAATAGGCTAGAGCCAAGTTAATTGCATTGTT
TCGTCTAAAGAGAAAAATGCGTATAGAGTAACGTTACAAAATAGCCTTATAAATTGGCTG
CTTAGCGCCAGCACACCCACACAGAAATACATACGCAGAAGGAATTGCGCGGCGAATGCC
ATCGGTTTGTCAAGGGCTGCTGTGATATAAGAAACAGGTTTTCTCTGTTTAGGTGAGAGG
TCCACAGAT

>LG29893_g1_i1
TTTTTTTTAAAAATCAAAACAAGAACATCTTTATTTAAAACATACAGGATTTACAAATCAGTACAAAATT
TCATCTTATAATATTTTCATTATTACCATATGATGTATTTACATACCAAAAGCATATTGAAATATATTCTT
TGCTGGCTGTTGTATGTACATAAGAATGTAATGCTAAGTAGAGTATTGCAGTTCCAATACTCATCTAC
CACTGACCAGTAATGCCACATCAGTAACTTAATTTGCATAGCTATTACTTTTATAATTCAGACATTTTC

ACTAAAATACTGACTAATAAGTACAAATCTAAATTACCATTTGTAGTAAGCTATCAAGTGTGATAAAAT
TCAAGTCTTTTTACATGTATCACACGTGACAAGATACTGCTCTAGGCACTGTAGTGGACTGTCACTTT
ACACTTGGCTTTTCAAAATTATGTCTTCCTATTACTAAATTAATTCTTTATAATTCTACATATAGATTACT
TATGTTTTGAAGTACAATCATTCCAAGTTGCCTTCTTCAATGAACCTAGAATACCACTTACCACTTTGC
TCAAGATGTTGAAAAAATAGACAACATATGTTGAAATAAATAGTTTTATTTGTATTTATCATTTAAAATG
GGAATACATATTTCTGTAATGCATTAGTGAAACCTGAAAATGGAAAACAGCAATAAGATACTGGAAAA
ATAAAGATGAACATCTGAAGAACACCAAAAGGAGCTACAGTCACAGCTAAAATGTTATCATTTGAGAT
CTCTCTGTAGAACCTACAAAGTCAGAGGTAACAGAGATGCTAGAATACCTGTGGACATGGGAGCTAA
CTTTTTTTTAAAAAGCTTAGCTGATATAAATAGAGAGTAGCTGATCTAGATAAAAAATAAAATGGAAAT
CTGGGACCTGGACATTATTTTATTTGATGTAGGTATGCAGTTGCACTTATATAATATCTACAGCTCCAC
TCTCATTATTCTGCAACTCACAGGTTTATAAATCATGGAATATGGTTTGGGTAGTGGTCTGGACTCCA
AATGTATAACTGGACTCCTCTAGATATGCAGTCGTGCATGTGATTTCTTCATTGTGTACTGGCCCCAT
AAGAATTATGATGGACTAGTGTATGCAGGTCAAACTGTGCACCAGCACTCTGCAGTTGTGCACAAGG
TTTATGAATAGCACTCATATACTATAACACATTCACTCTAAAAGAAAGTGGCAGTGCAGTCATTAAGAA
ATAAATGCTACATTTCATAAATTACTGTTAAAAGTTTTAGGCTTCCTGGCCAATCATTAATTAAGGATTA
TATAATACTAGCTAGGATCTCAGTCTACATTCTAACCTTAGGGGACACCTCTGTATTATATAGGCATG
GTCTGTCTGACACATTCACCTAATGTCTACTGCTATATTCATACTGTGTACTTTTTAATTTGAAATAAAT
ATTACATGGAAGGTGCATCTGATTTTATTTACTTGAGAACCTCTAGACTGTTAACATTTTTCTCTCTGC
TCCAGTGATCTCCAAAATATCTGACTTTGATAATGCTGTTGACCAAAAAAGTTAACCACTGTAAACTCT
GGTGCCCATTTAGCATCAGTGACAATATGCTATGACGATGTTATCAGACTTAGTGCAACTACTTAAGA
CACACTGCTTATAAACATCACTTGAGTGGATGATTATAGATGCAAAAGTCAGAGCTTCATAATTTGGC
TAACCAACCCAGAAATGATGTACAAACCTAATACTGTTCATCGTCATGAGGTAGTAAAACAAAACAAA
TCCACTTTTGGGACAATAGCTACTATACGTATAGGACATTTTGCTACAGTGCCTGCAAAGCTTTGTAT
GTACTTTTTTGTGGCATGTATAATCTTGAAGAAGTCACATAATAATTTACAACACATTTAATTTTTTTCT
GAAATACTGATGCAAAAGCGTCTTGTATATCAGTTGTAGACCGTTCCCTTTGGAAGACTGAAGATCTG
TAGTCCATGTTGATTAAGGCAATGTGCTGTTCATCCCAAATATTTGAACATCTTTTTTTCACAAGTAGC
TCCAGTGTACCCAGGACGGCATGAACACACATTTGGAAGTATACATCTGCTCCCATACAGACATTTTT
GTTGACATATAGGTATTTGGCATTGTAGTCCTTCCCAGCTGTTTTGACAGTGGCAGGTGTTAGGTCCT
ATACATTCACCTTCATTCTTGCATATCTGAAGACAAATAGGTATGCTACACCTTTTTCCTTTCCAGCCA
GATGGACAAGAGCAAATGTTTGGTCCAACACACTTTCCTCCATTCATACAGATGGGATCGCAAACAC
TTTTCTGACACCTCTTTCCAGTGTAACCATCAGGGCAAGTGCACACGTTGTTTCTCATGCAGTGGCCA
CCATTTTTGCATGGTGGACTACATATGGCATTCTGGCACTGTATACCAAAGAATCCATTTGGGCAAAG
GCAAACATTTTGTTTAATGCATGTTCCTCCATTCAGACACACTGGGCTACATACAGCTGTACTACAAG
TAGGTCCAGACCAGCCTGCTTTACATTTGCAAACATCTGGAGTAATGCATTCACCTCCATTTTCACAA
TGCCGATTACAAACCATGGTTTCACATCTTGGCCCCACAAAACCATAAGGGCAAGTACAGAGGTTTC
TTGCCAGACAAGTACCTCCATGTTCACAAGGTGGATCACAGTGAGCTGTATCACATGTTGGTCCATC
ATATCCAGAAGGACACTTGCAGACATTTGGTTCGATACATTTTCCAAGGTTCTTACAATCGGGTCGAC
ACACAGCAGTTAGGCAGTTATATCCTGAGTACCCAGGTTTGCATCTGCATATATTTGGTGCTGTGCAT
TCCATATTCTTGCCACATGGGTGCCTGCAAAATGCTTTGCATGTGATTCCATCACCATAGTACCCAAA
TGGGCACCTTCCACACTTGAATGTACCATTTAAAGTTGGCTCACAGGGAACACCAGGGAAGCAAGGC
GCATCCGCACATGTAATTATCTTTTGGACTGATGTATTCATCACATCTCCAACTTTACCCACCGCTGA
CTGTTCCACTGTACCAGGAAGCACAGACTGCTCATATGTAATGCTATGTGTGACTGATGCACTTTCAG
ATCCGTTTCTTGTTGTGCTTCCAATCACAAAACCATTTCTGTCATTTGTAGATGTTGTTTTCAAGTTTTT
TAAATTGCTTTCATCACCTGTTTGCAGCAACACAGTTCTCCTGGTTGGTAAAGGCCTGATCCCCAGAG
AGGAAGCATTCCATTTCATTTGTGTTACTGGAAATCTTCTTCTTGAATCTCTGTTGTCTTGTGCAGTGG
AAGTATGAACTGGTGATCTACCACCAGAGGAATGAATAGTGATAGTACCCTTCTGAGGAACTAATGG
CTTCCTGCTTGCTGTAGCAGTAGCAGTGGTTGTAGCAACAGTGGTAGAGGTAGTAGCTGTGGCAGA
GGAAGTAGTAGTTGCAATAGCAGCAACAGTCATAGTAGTATTTGGTGGAATACTGGTGGCTGTTTTA
GTAATACTGGAAGTAGTGGTAGTGACATCAGCAGGAAGGGTACTTTGTTTTGCAGAAGATGCTCTAA
AAAATGGCAGTTGTGTGGTATATAGTGTTTGCTGTACACCAGAACCAGATGTAGCTGTAGAGTGGCT
GATATGCAATTGAGTAATCTGACTTTCATCCAAAATAACATAATCTTCTTTATAACCTTGTTTTGTATTA
ATCTCTTCATCCTCATCCTCATAATAAATGTAATCGTCTTTAAAGTTTTCAGCAACAACAGACAAAGAA
GGGACTCCATCAGTGGGTTGTTCAATGCATATCTTTCCATCTCCTTGCATTCCTGCAGGACAAGAAC
CACACTGATAGGATCCAAATGTGTTGGTGCAGATCACTCCAGCAAAACAAGGATTTCTCTTACACTCG
TCAATGTCCTCTTGACAATTGTTGCCTCTCAGCCCAGTTGTGCACTCACACGAGTAGCTGTTTATTCC
ATTAATGCACCTTCCTGAGCCACAGGGATTAGAACTGCAGACATCAGTAATTACTTGACAAAACTGCC

CTTCAAAGCCAAGGGGACACACACAGAGGTACTCTCCACTTCCAGGTGGCCAGTTAATATTTGTAAC
ACACGATCCTCCATTTTGGCAGTCACATGGTTTGACAGCAACCTCTACAGTTATTCTGGTCTTAGCAT
TGCAGTCATCTGTAATAGAGAACACAAATGACTCTGGGAGTTGTAATGTCACTTTCCATATAAGCAGT
CCAGCAGGAGAAAGACTGGCACCTTCAGGTGCAGAGTCCAGAGTGAATACTACCGAGGAACCTTCT
GGGTCTGTTGCTGTGAACTGGTATACAAAGTTTTCCCCATAAAAGGTTTGAAGCTTTTTGTGTGGTGT
CTGAAGCACTGGTGGATGGTTGTTTTCAGTAACTGACCATGTAAATTTGGATAGATCAGGCTGACAC
ACCAGGCAAGGAGTTGTGGGATTTGTATCTCCTTTCCCATAACACAGTCCTTCTATGTTGCAGGTTTC
CTCCTTCAGACTACACAGTCCATTAGAATGTGGGTCACAGATCTGACAAGCACCATCATATAAAGTCA
ATATTTTGGAGTTGCTGTATAAGTAGCCATCATTTGAAACCTTTACTTCCCATCTTGCAATTGGTTTGT
CATCAATTAGATCCATGCTGTCAGGTGACTGGACATTTCCAGATGGAAGCTGACATTCCATGGCTCT
AGTGCTAAGGAAAGTAGCCACAGTTGTCTGAGGGTTTCCAGGGATCCATTCTCCATTGCTATACTGT
TGTTGTGTCACTTCACATTTAAGGCTAGGGGAGTCTCTGAATCCTTCTCCAAAAACTCGTACCAAAAT
GCAGTCATAGAGACGGACATCACACAATCCACTGTTTTCTAGTTCTGTAATTTCTGGCAGCACATCAG
TCAGAATGCTGCAATCATAAGAGCCAAAGCCTGGGAAACATGCACAGCCCCATTCTGCACACTCTCC
ATTTCCATTGCACAGATTAGGGCACCTCAGCACTGAAATAATCTCATCAGCCAGTACACCTTGTTCTT
TTGTAGTAGAATAGCCATCTTCTAGAACTTTCCTTTCACACTCATTTTCTAATAGAGCCAGGCCAGCTT
CTATCCAGCTGGGATCATCCTTCAGTAACAAGTCTAAGACACACATGTCAATTACATCCAAGATTCGT
ATGCTAAGTAGGTCTGCACACAGGCTTCCTATACTGGAATTAGTAATAGTTTGCTGGCACAGTTCTTG
GGCCTTGTATTCTGTAAGGCCAGAAGGTGTGGGCCAAGTGGGTAAGGCTTCTTGATAGCCCTCTGTA
GTGTGGTCTTCTGGGAAAAAATAGTTGAACCCCTCAAGATCAACTTGACTTAGACTCTGAAAGGGAA
ATGTTGGAAGGTATTCATAATAACCCTGACGTTTATGTCTGTATCTTTCAGAGACAGGTTTCAGTGGA
ATATCACTTTGCTTTTGAAACCACCTTGTTTCTTGTTTCAGTCCTTCATATTTTTCCTTTGAGTTTGTTC
TTTTAGAAAGTGAAATTGAGGTGTAATGACCTTGTGCAACTCCTGTATATTCGGTTTGATTGCTCATTT
GGCCACGTGCAAAAGGAGAAACAGAAAAGTCTTTTTGAATTAGTGTATCACGTTTACTTAGCTCTCTG
GTGAGCTCAAAAGAATTGACATACTCTGCTGTGATATCTAGGACTGGTATTAGTGCTGTATGTTGGAC
ATCTTCTTTGCCTGAACAAATGGATAAATGTCCAGACTTCTGGGAAGTATCAGGCTTGCTAGAAGATT
TTTGTGACTGGGCAATATCTTCACTGCAAGTACAGTACGGTCTTCTTCTTGTTGAGGTGCTTGATGAT
GGTACTTTGTCAAACAAACTGCTACCTGGTACAAGTCTCCACTCATCCACAAATGCTTGAGCATTTTC
CAGGATAGTGATCCCATCAGCACTCTGGAAATCATTGTCAGGGTCTTCATCAAAGGAACCACATAGT
CCCTGAGTATTCCTGTAGTCTGAGCTAGGTGCCCTAAGTGACAAGCTCATTCCCCATTCACTGACAT
CTGCACGGATAAATGCTCCTGAAGAAAATGATACTGTTACTTTTCTTCCTTGGTAGGATTCTGTGATT
CGTACATTACCTCCTGTGATGTCACGACTTTTCACTGATAAATGCGGTCTAGTCTCATGTAGTTGGCC
ATTGCACATGTCAAAAGAAATTATGTCACCTCTTTCTCTGGCAACAAAACCACAAGTACAAGCAGCTT
GGTAATGGAGGCTGCCACAGTCCCACTGGCGAACATGAACTTCAAAATCGCGATTTGTACTCTTGTA
TAAAACAAATGTTCCAGTTTTAAAATTATCAAACCTCCTGCCATCAAATGTAATAATATGTGGGTCTGT
GAATGAGTAGCAGTATGCAGTAGGAACATCCTTTACTGTAATCTGTATACCCTCTGGGGCATAGCCA
TTCCAAAGAAAATTCTCATGAAGTATAGGATGAGCCACAATTTCTGTCACTTTGTCTCCATCATGTGAA
AAATCTGTCACAGCTGTGAAGTAAACAGTTCCACTTCCACAGATCTTATTATGGCAAGGTAACTGGTG
CAAGTCCACTTGACAAGATGAAAGGGCAAGGTCTGCAAATGGATTCTGTTCATCTTCTTTATTCACTG
TCTTTAGCTGGAGAGAAATCTTGCAGTTATGGTCTTGCTGGCTAGTGTCAGAGCACGGGATGGGTAC
TGAGCTCTCAATTGTCAGCCTATGTTCCATTCCATCTTCTGATATTGTATGAACCTCAGGGTATACCTT
TACACCAGCAAAAAACTCCTTGCTTTCAATGGAAGGGCTGCGCACATCAGGGGTCTCCAAGAAGAAA
CTGGAACTGGAGCAATAAATCTTGTCTCCAAGCCTGAGATTTATGCCATCCAGTTCTATAAATGAGAA
TGTTTGGACTGTCGTTTCCTGCCGCAGTTCTTCCTTGACTCCATAAGGGGATAATCTGGACCAAGTAA
CAACAAACCCAAGAGAGCTGTTTGTGCCTGATACATCAAATTCACACTTTACGTAAATAACACCCCCT
GTCAACTCTGCCACAACTTCTGGAGTAGACGGTGATGGTGGATATTTAGCTCTGCAGGCTCCTCCAG
CTTCTGTTTCTTCAGGGCCACAAGACTGCAATTTTAAGCCAGAAACTACTTCAGCACAGTATCCCATA
CATCCCTGTGTTGGTTGGAGCAAATACACAAAGAAATCTCCACAGTTTCGCATACTAATTGGTATTCG
GAAAAGACAGCAGTCCTTAGTAGCACTGAAAAAAAACTGCCATGTTGCACAGGCTGTTAGCTGCTTG
ATTTCTCCGGGCTGTGGAAGAGATTCAGAGTCCTTCAGTGACAGCCATATAGGGGCTTGAGTTCCAC
AGTGATTCATCTCAACACATTTGGTTGGCATCTCTGCTGGTTTATCAAAGATCTGGAAGCGGTACCAT
CCTGCAGTTAATGAGTGGTCACATATCAAATCCTGAATAGCTGATTGTTGGAGTCTTAAGGAATCAAA
ATCAATGCTTCTATATGGATTTTGAAGTATTTGATGTCCCCCAGGAGAACATTCATGTGCTTGCTGAC
AGACAGCTATTTTTAAGTGCATTACTGAGATCCAGATAATAAAACAATCTGCAAGGAACTTTTGAGTC
ATGGCTAACAAAATGTATTATTTCATTCCCAGATGCTTGATACTTCTGTTTTTGCCTCTTTACAGCTTA

ACTTAAAACAAAAATGTTCCCAGTACCAATTGACCTGTCACATTACTATGCACAGTTATCAATGGCCA
GTGGACTCAAGTGACTGCACTACTAGACACTCTGTAATCCTACACATGATGCTG
