## Supplementary File 2 for "von Willebrand Factor D and EGF Domains is an evolutionarily conserved and required feature of blastemas capable of multi-tissue appendage regeneration"

>pCAG_td_tomato_MO1_binding_stop
ATGGTGAGCAAGGGCGAGGAGGTCATCAAAGAGTTCATGCGCTTCAAGGTGCGCAT
GGAGGGCTCCATGAACGGCCACGAGTTCGAGATCGAGGGCGAGGGCGAGGGCCGC
CCCTACGAGGGCACCCAGACCGCCAAGCTGAAGGTGACCAAGGGCGGCCCCCTGCC
CTTCGCCTGGGACATCCTGTCCCCCCAGTTCATGTACGGCTCCAAGGCGTACGTGAA
GCACCCCGCCGACATCCCCGATTACAAGAAGCTGTCCTTCCCCGAGGGCTTCAAGTG
GGAGCGCGTGATGAACTTCGAGGACGGCGGTCTGGTGACCGTGACCCAGGACTCCT
CCCTGCAGGACGGCACGCTGATCTACAAGGTGAAGATGCGCGGCACCAACTTCCCC
CCCGACGGCCCCGTAATGCAGAAGAAGACCATGGGCTGGGAGGCCTCCACCGAGCG
CCTGTACCCCCGCGACGGCGTGCTGAAGGGCGAGATCCACCAGGCCCTGAAGCTGA
AGGACGGCGGCCACTACCTGGTGGAGTTCAAGACCATCTACATGGCCAAGAAGCCC
GTGCAACTGCCCGGCTACTACTACGTGGACACCAAGCTGGACATCACCTCCCACAAC
GAGGACTACACCATCGTGGAACAGTACGAGCGCTCCGAGGGCCGCCACCACCTGTT
CCTGGGGCATGGCACCGGCAGCACCGGCAGCGGCAGCTCCGGCACCGCCTCCTCCG
AGGACAACAACATGGCCGTCATCAAAGAGTTCATGCGCTTCAAGGTGCGCATGGAG
GGCTCCATGAACGGCCACGAGTTCGAGATCGAGGGCGAGGGCGAGGGCCGCCCCTA
CGAGGGCACCCAGACCGCCAAGCTGAAGGTGACCAAGGGCGGCCCCCTGCCCTTCG
CCTGGGACATCCTGTCCCCCCAGTTCATGTACGGCTCCAAGGCGTACGTGAAGCACC
CCGCCGACATCCCCGATTACAAGAAGCTGTCCTTCCCCGAGGGCTTCAAGTGGGAGC
GCGTGATGAACTTCGAGGACGGCGGTCTGGTGACCGTGACCCAGGACTCCTCCCTGC
AGGACGGCACGCTGATCTACAAGGTGAAGATGCGCGGCACCAACTTCCCCCCCGAC
GGCCCCGTAATGCAGAAGAAGACCATGGGCTGGGAGGCCTCCACCGAGCGCCTGTA
CCCCCGCGACGGCGTGCTGAAGGGCGAGATCCACCAGGCCCTGAAGCTGAAGGACG
GCGGCCACTACCTGGTGGAGTTCAAGACCATCTACATGGCCAAGAAGCCCGTGCAA
CTGCCCGGCTACTACTACGTGGACACCAAGCTGGACATCACCTCCCACAACGAGGA
CTACACCATCGTGGAACAGTACGAGCGCTCCGAGGGCCGCCACCACCTGTTCCTGTA
CGGCATGGACGAGCTGTACAAGGCCAACGCAAGGATGTATGGGATATTAG

>pCAG_MO2_binding_kozak_td_tomato_stop

ACTGTGAGGAACTGTGATGGTTTCTGCCACCATGGTGAGCAAGGGCGAGGAGGTCA
TCAAAGAGTTCATGCGCTTCAAGGTGCGCATGGAGGGCTCCATGAACGGCCACGAG
TTCGAGATCGAGGGCGAGGGCGAGGGCCGCCCCTACGAGGGCACCCAGACCGCCAA
GCTGAAGGTGACCAAGGGCGGCCCCCTGCCCTTCGCCTGGGACATCCTGTCCCCCCA
GTTCATGTACGGCTCCAAGGCGTACGTGAAGCACCCCGCCGACATCCCCGATTACAA
GAAGCTGTCCTTCCCCGAGGGCTTCAAGTGGGAGCGCGTGATGAACTTCGAGGACG
GCGGTCTGGTGACCGTGACCCAGGACTCCTCCCTGCAGGACGGCACGCTGATCTACA
AGGTGAAGATGCGCGGCACCAACTTCCCCCCCGACGGCCCCGTAATGCAGAAGAAG
ACCATGGGCTGGGAGGCCTCCACCGAGCGCCTGTACCCCCGCGACGGCGTGCTGAA
GGGCGAGATCCACCAGGCCCTGAAGCTGAAGGACGGCGGCCACTACCTGGTGGAGT
TCAAGACCATCTACATGGCCAAGAAGCCCGTGCAACTGCCCGGCTACTACTACGTG
GACACCAAGCTGGACATCACCTCCCACAACGAGGACTACACCATCGTGGAACAGTA
CGAGCGCTCCGAGGGCCGCCACCACCTGTTCCTGGGGCATGGCACCGGCAGCACCG
GCAGCGGCAGCTCCGGCACCGCCTCCTCCGAGGACAACAACATGGCCGTCATCAAA

GAGTTCATGCGCTTCAAGGTGCGCATGGAGGGCTCCATGAACGGCCACGAGTTCGA
GATCGAGGGCGAGGGCGAGGGCCGCCCCTACGAGGGCACCCAGACCGCCAAGCTG
AAGGTGACCAAGGGCGGCCCCCTGCCCTTCGCCTGGGACATCCTGTCCCCCCAGTTC
ATGTACGGCTCCAAGGCGTACGTGAAGCACCCCGCCGACATCCCCGATTACAAGAA
GCTGTCCTTCCCCGAGGGCTTCAAGTGGGAGCGCGTGATGAACTTCGAGGACGGCG
GTCTGGTGACCGTGACCCAGGACTCCTCCCTGCAGGACGGCACGCTGATCTACAAG
GTGAAGATGCGCGGCACCAACTTCCCCCCCGACGGCCCCGTAATGCAGAAGAAGAC
CATGGGCTGGGAGGCCTCCACCGAGCGCCTGTACCCCCGCGACGGCGTGCTGAAGG
GCGAGATCCACCAGGCCCTGAAGCTGAAGGACGGCGGCCACTACCTGGTGGAGTTC
AAGACCATCTACATGGCCAAGAAGCCCGTGCAACTGCCCGGCTACTACTACGTGGA
CACCAAGCTGGACATCACCTCCCACAACGAGGACTACACCATCGTGGAACAGTACG
AGCGCTCCGAGGGCCGCCACCACCTGTTCCTGTACGGCATGGACGAGCTGTACAAG
TAG
