## Supplementary Figures for "von Willebrand Factor D and EGF Domains is an evolutionarily conserved and required feature of blastemas capable of multi-tissue appendage regeneration"

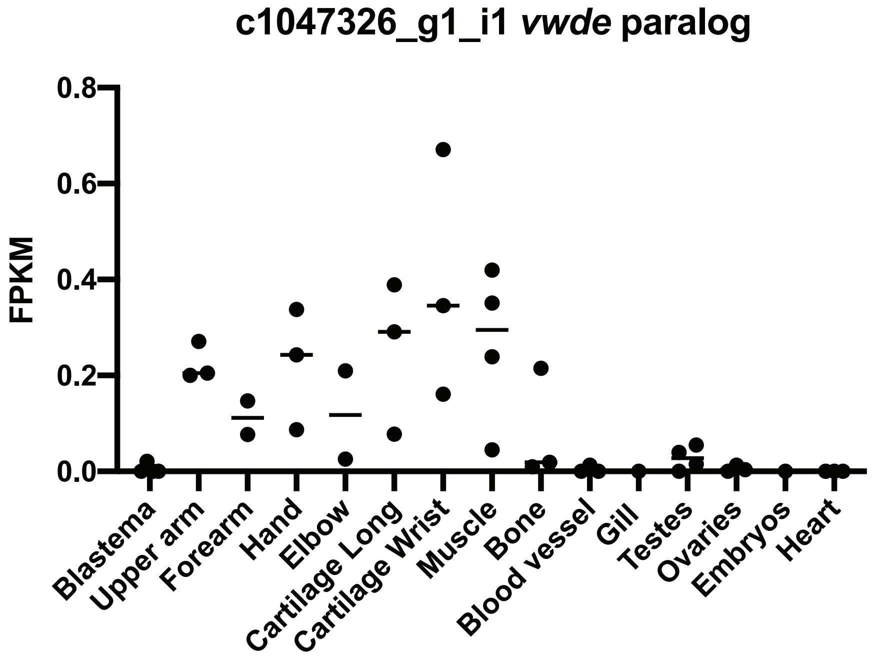


**Supplementary Figure 1. Axoltol paralog to *vwde* is not regeneration-enriched.** Expression profile of *vwde* paralog contig number c1047326_g1_i1. FPKM values across tissues samples mined from Bryant et al. 2017 Cell Reports. Line indicates median value.

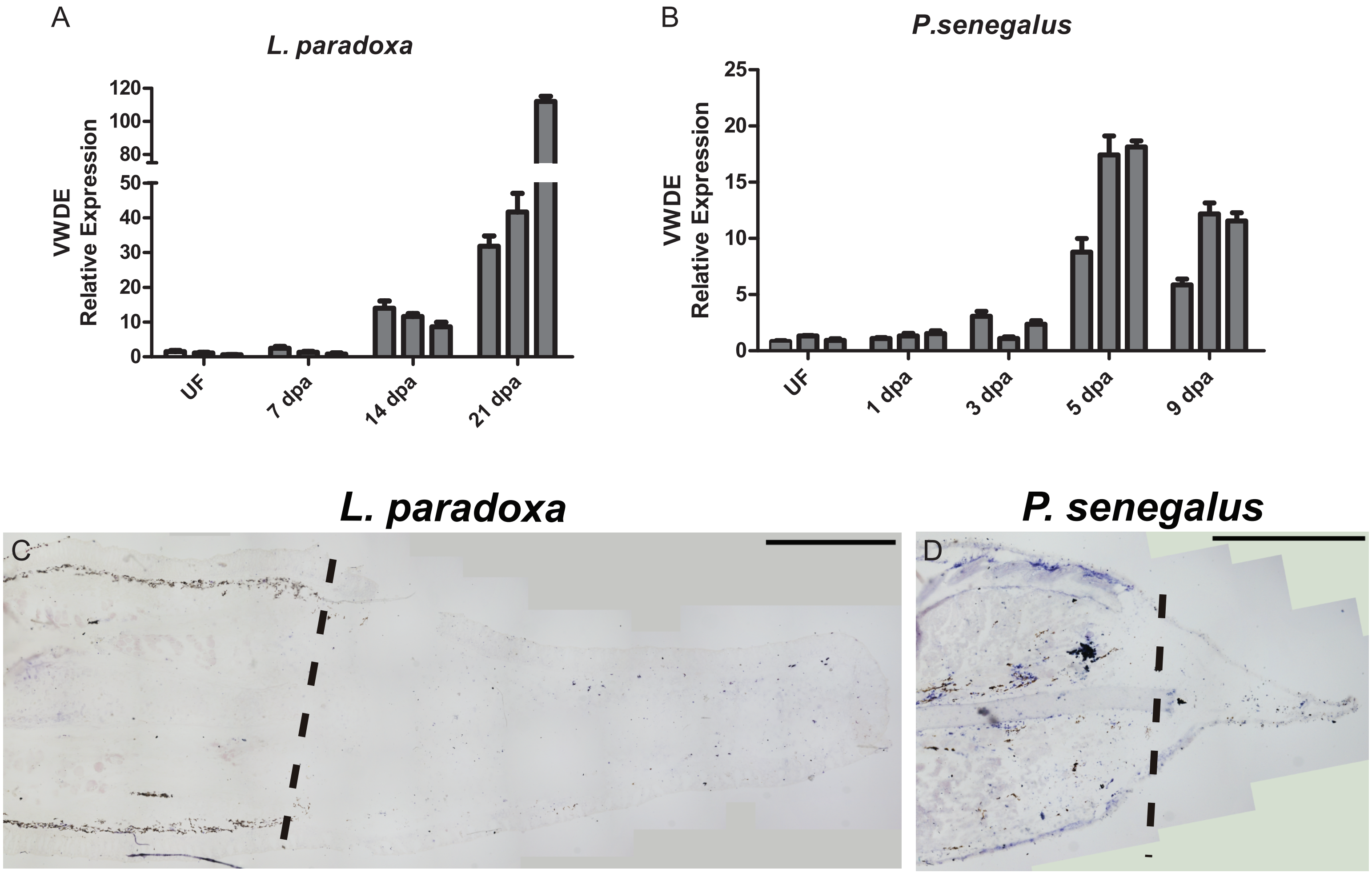


**Supplmental Figure 2: vwde is blastema-enriched in blastemas of other species.** Expression of *vwde* during fin regeneration in *L. paradoxa* (A) and *P. senegalus* (B). qRT-PCR data for *vwde* in uninjured fin (UF) and regenerating fins at the specified number of days post-amputation (dpa). Relative expression was calculated using *sdha* (*P. senegalus*) or *polrc1* (*L. paradoxa*) genes as endogenous control and the mean value of the normalized Cts of all three UF samples as reference sample. (C-D) In situ hybridizations using sense probes for *vwde* in *Lepidosiren paradoxa* and *Polypterus senegalus* pectoral fin blastemas. Longitudinal histological sections of fins from *L. paradoxa* at 21 dpa (C), and from *P. senegalus* at 5 dpa (D). Dotted lines indicate amputation site (Scale bars, 1 mm in all panels).

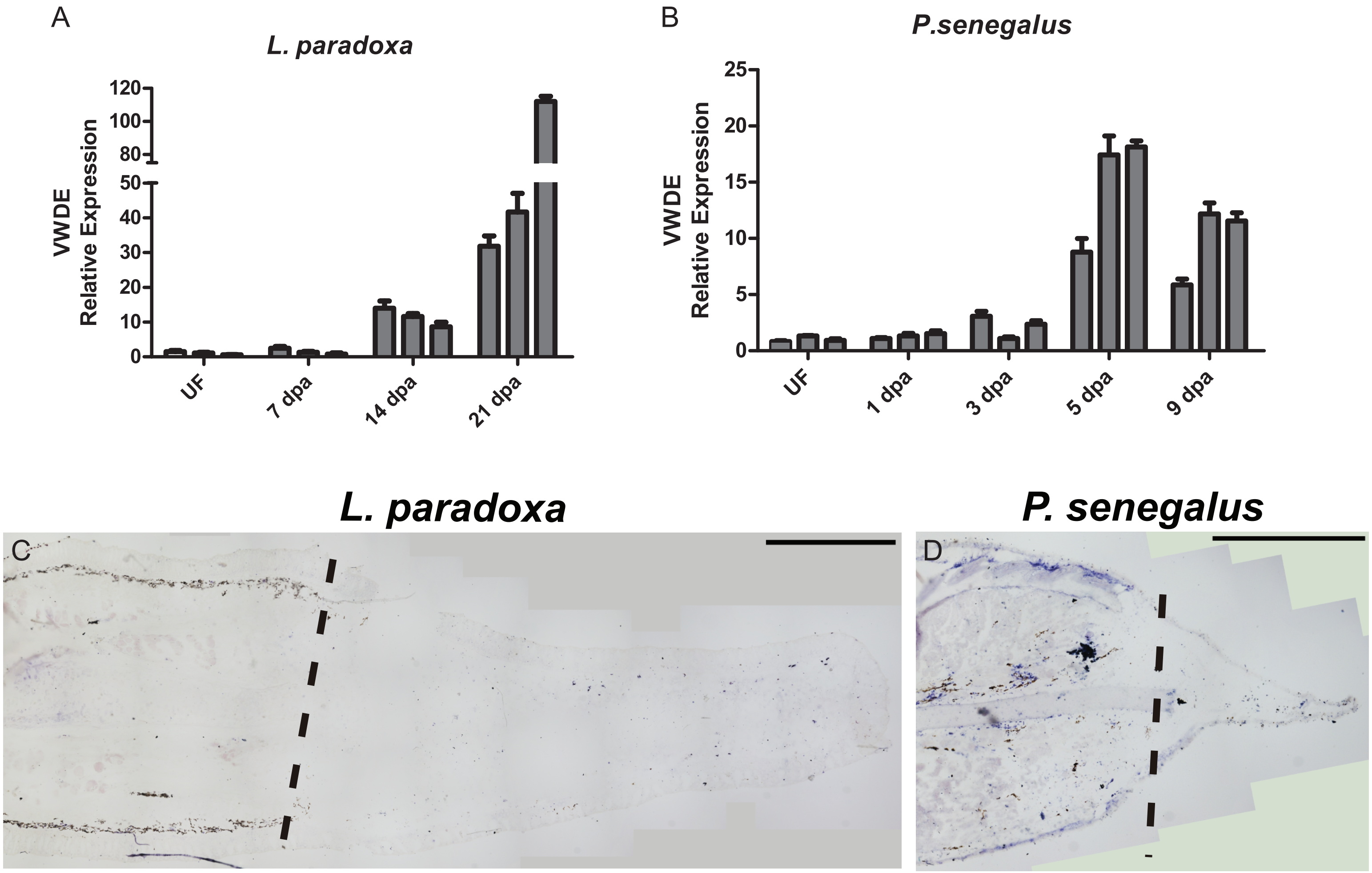


**Supplmental Figure 2: vwde is blastema-enriched in blastemas of other species.** Expression of *vwde* during fin regeneration in *L. paradoxa* (A) and *P. senegalus* (B). qRT-PCR data for *vwde* in uninjured fin (UF) and regenerating fins at the specified number of days post-amputation (dpa). Relative expression was calculated using *sdha* (*P. senegalus*) or *polrc1* (*L. paradoxa*) genes as endogenous control and the mean value of the normalized Cts of all three UF samples as reference sample. (C-D) In situ hybridizations using sense probes for *vwde* in *Lepidosiren paradoxa* and *Polypterus senegalus* pectoral fin blastemas. Longitudinal histological sections of fins from *L. paradoxa* at 21 dpa (C), and from *P. senegalus* at 5 dpa (D). Dotted lines indicate amputation site (Scale bars, 1 mm in all panels).

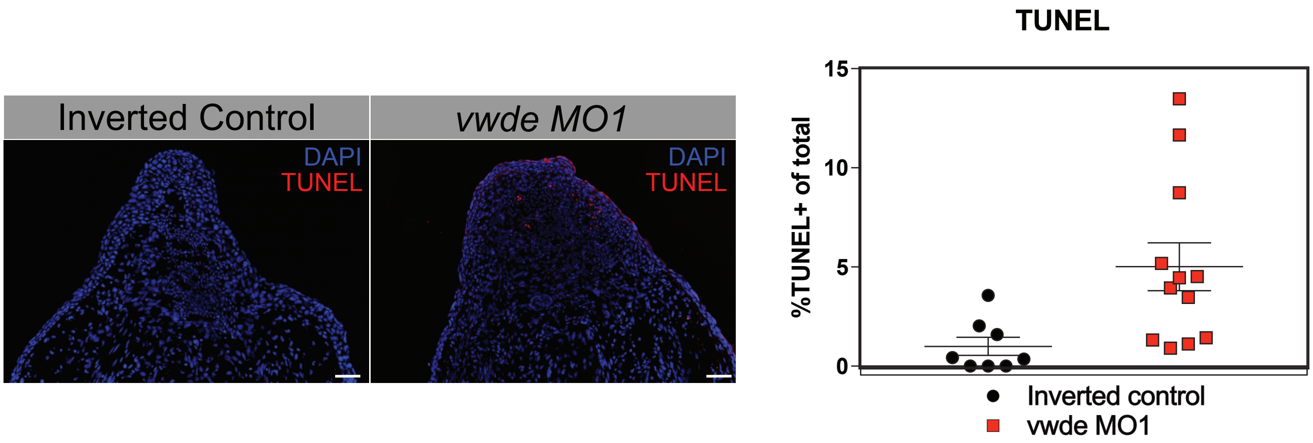


**Supplemental Figure 4: TUNEL staining after VWDE knockdown.** (A) Representative TUNEL stained sections of blastemas 10 dpa (3 days post-electroporation) of control morpholino (VWDE morpholino 1 inverted) and VWDE-targeting morpholino 1. Scale bars are 100um. (B) Quantification of percent of blastema cells positive for TUNEL in control (*vwde* MO1 INV) and knockdown (VWDE MO1). Each dot represents a limb, 3-4 animals per group. No difference between treatments detected via nested t test.

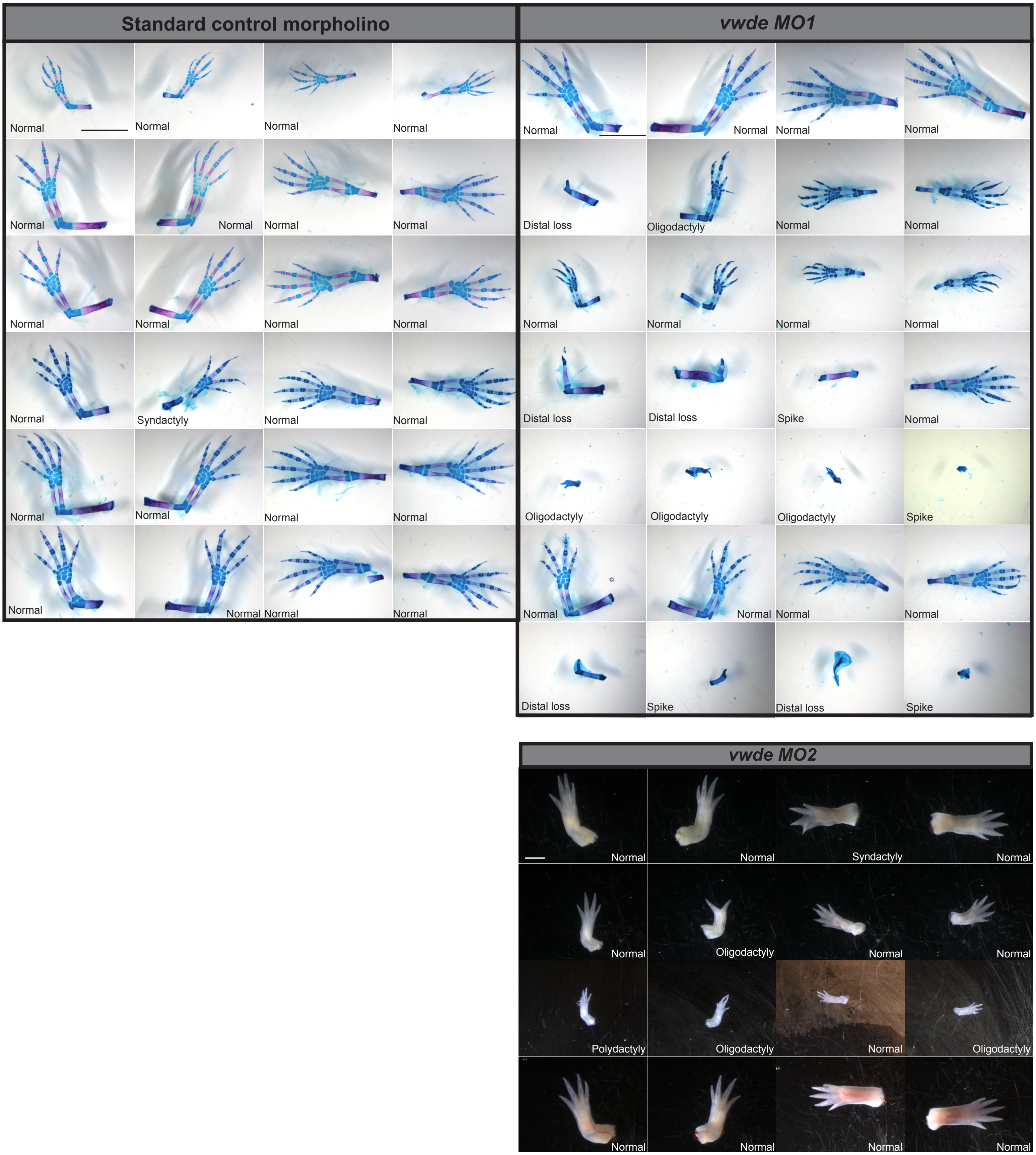


**Supplementary Figure 5: Knockdown of VWDE impairs regeneration.** Alizarin red and alcian blue skeletal preperations following the full course of regeneration (10 weeks, top) or limbs. Control (*vwde* MO1 inverted, left panel) and VWDE knockdown (*vwde* MO1, right panel). Each row of each treatment is an individual animal, all four limbs are shown. Bottom left labels indicate phenotype observed, see Methods for description of scoring. Scale bar is 5mm.

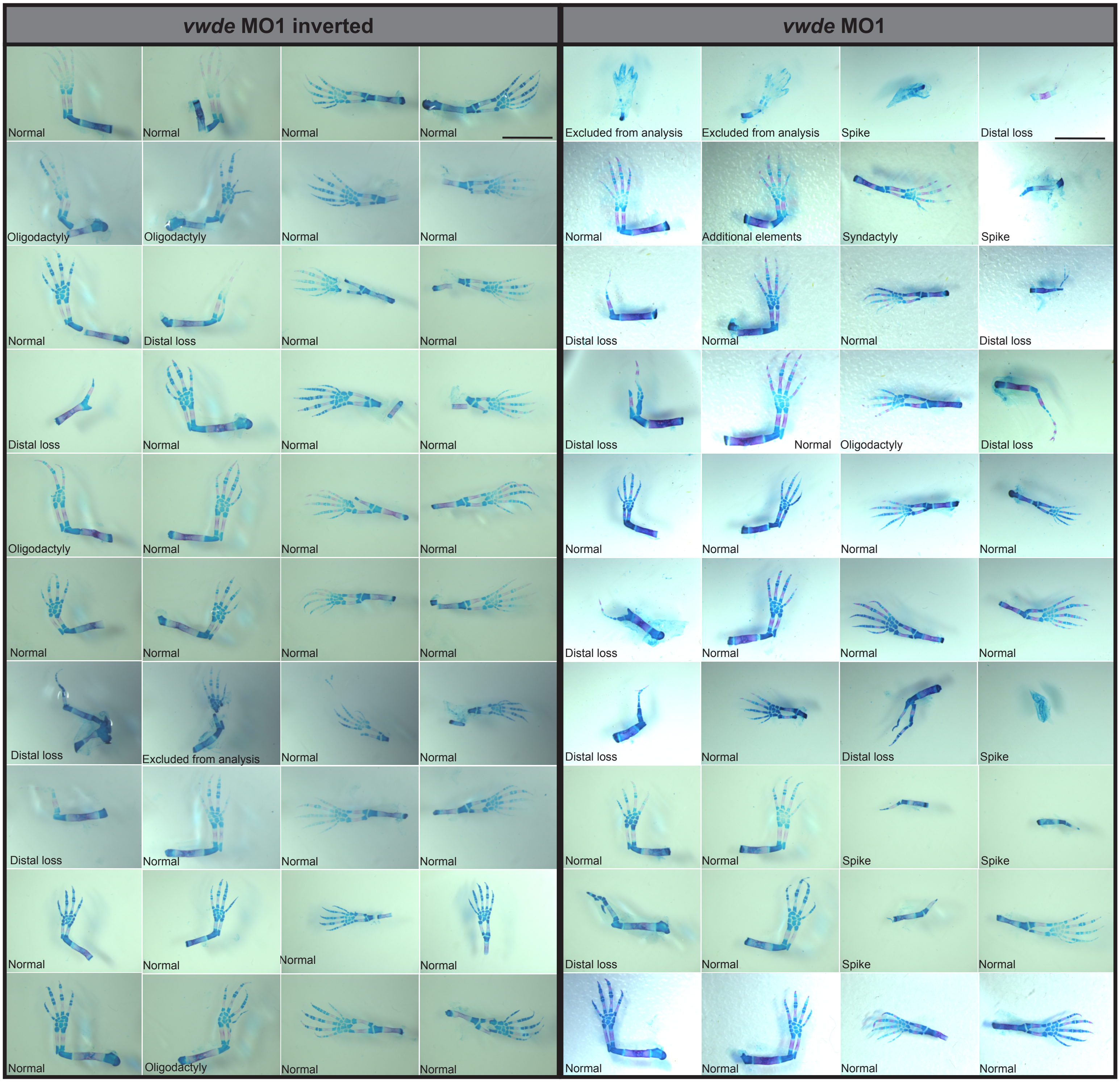


**Supplementary Figure 6: Knockdown of VWDE impairs regeneration.** Alizarin red and alcian blue skeletal preperations following the fullcourse of regeneration (10 weeks). Control (*vwde* MO1 inverted, left panel) and VWDE knockdown (*vwde* MO1, right panel). Each row of each treatment is an individual animal, all four limbs are shown. Bottom left labels indicate phenotype observed, see Methods for description of scoring. All images taken at the same magnification, scale bar is 5mm.
